# Astrocytes in the Dentate Gyrus Shift the Equilibrium from Pattern Separation to Pattern Completion and Inhibit Fear Extinction

**DOI:** 10.64898/2026.10.05.755780

**Authors:** Muxiao Wang, Maltesh Kambali, Rajasekar Nagarajan, Jinrui Lyu, Hsun-Yi Kuan, Xinzhu Yu, Uwe Rudolph

## Abstract

Pattern separation and pattern completion are core hippocampal computations linked to memory interference and overgeneralization in many neuropsychiatric disorders and to mnemonic discrimination deficits in neurodegenerative conditions. Astrocytes are increasingly recognized as important integrators and modulators of neuronal activity as many studies have shown changes in animal behavior and neural circuits induced by manipulation of astrocytic activity. The role of astrocytes in the dentate gyrus for learning and memory is, however, largely unknown. To investigate the functional significance of astrocytic activity in the dentate gyrus, we expressed the calcium extruding pump hPMCA2w/b (CalEx), resulting in inhibition of astrocytic calcium activity, the G_q_-coupled receptor hM3Dq, or the G_i_-coupled receptor hM4Di, allowing chronic activation of G_q_- and G_i_-dependent pathways by a designer drug, in dentate gyrus astrocytes. Our main finding is that inhibition of astrocytic calcium activity by CalEx resulted in improved pattern separation and impaired pattern completion. In vivo calcium imaging revealed that CalEx expression in astrocytes altered the neuronal activity pattern in dentate gyrus neurons with an overall inhibitory effect. This suggests that in the dentate gyrus astrocytes promote the activity of these neurons. Furthermore, chronic activation of hM4Di resulted in impaired between-session fear extinction while chronic activation of hM3Dq resulted in impaired within-session fear extinction and improved pattern separation learning over multiple days, suggesting that activation of astrocytic activity impairs mechanistically separable forms of fear extinction in a G_q_- or G_i_-pathway-specific manner. Our findings reveal that dentate gyrus astrocytic calcium shifts the balance between pattern separation and pattern completion towards pattern completion and thus modulates an essential neuronal functional output of the dentate gyrus. Chronic activation of astrocytes inhibits fear extinction, and thus also affects management of memory interference. Our findings imply that astrocytic modulation may be beneficial to treat cognitive deficits in disorders in which overgeneralization is a problem.

## Introduction

Traditionally astrocytes have been regarded as secondary support cells for neurons. Although astrocytic calcium signaling was discovered more than three decades ago ^1,2^, it is only in the recent decade that astrocytes have been increasingly recognized as having important roles in regulating the function of neuronal circuits and the behaviors they facilitate ^3–5^. Thanks to advances in the tools available to investigate astrocytic function ^6^, our understanding of neural circuits beyond neuronal components has been substantially improved. These recently developed tools enable studying the function of astrocytes by manipulating the activity of astrocytes via different pathways. As recent studies expanded our knowledge about astrocytic calcium-dependent signaling as a fundamental cellular mechanism underlying the participation of astrocytes in neuronal networks^4,7^, understanding its implications for animal behavior, especially cognitive function, becomes critical ^8^. Although many studies have investigated the behavioral consequences of manipulating astrocytic activity ^9^, our knowledge of the functional role of astrocytes is still limited. Generally, astrocytes are thought to modulate the function of the neurons and neuronal circuits that they interact with. However, modulation of the activity of astrocytes and neurons in the same brain region may have diverging or even opposite functional consequences. For example, activation of astrocytes in CA1 induces de novo NMDA-dependent long-term potentiation in CA1, resulting in memory enhancement, while activation of principal neurons in the same brain region resulted in impairment of memory^10^. While it is known that astrocytes play integral roles in the regulation of neural networks^11^, it is possible that astrocytes in a specific brain region might modulate some but not all behavioral functions that the neurons in this brain region are known to mediate. To gain a deeper understanding of how astrocytes modulate neuronal networks and consequentially behavioral output, their role in individual circuit locations needs to be investigated with multiple tools that activate or inhibit astrocytes via different pathways^6^. Moreover, the finding of a high level of diversity among astrocytes further complicates a global understanding of astrocyte function^12,13^.

In recent years, studies have shown that chemogenetic activation of astrocytic G protein-coupled receptor (GPCR) pathways in the hippocampal subregion CA1 affects neuronal plasticity and subsequently behavioral phenotypes such as memory formation ^10,14,15^. Chronic activation of the G_q_-mediated pathways in ventral hippocampal neurons and astrocytes has been shown to differentially affect memory and behavior^16^. So far, most studies to elucidate astrocytes have used tools that activate biochemical pathways in astrocytes (e.g., activating G_q_-dependent pathways with a G_q_ DREADD or activating G_i_-dependent pathways with a G_i_ DREADD), while a much smaller number of studies have investigated the effects of inhibiting astrocytes. It has recently been shown that expression of hPMCA2w/b, a calcium extrusion pump, also known as “CalEx”, in astrocytes leads to reduction of Ca^2+^ signaling and thus inhibition of calcium-dependent astrocytic activities^17^. By actively extruding calcium, this tool provides a chronic “knockdown” effect on calcium signaling which is commonly associated with astrocytic activity ^7,8^. When CalEx was expressed in CA1 astrocytes, spatial memory was impaired and contextual fear increased^18,19^. While recent studies using DREADDs and CalEx have advanced our understanding of the functional significance of astrocytes in the hippocampal CA1 subregion^10,14,15,18,19^, there is a gap in knowledge regarding the functional roles of astrocytes in other subregions of the hippocampus, such as the dentate gyrus (DG).

The dentate gyrus is part of the trisynaptic hippocampal circuit and has been known to play critical roles in many behavioral tasks including anxiety-like behaviors ^20^, reversal learning in the Morris water maze (MWM) ^21^, latent inhibition of conditioned freezing ^21^, fear extinction ^21,22^, pattern separation ^21,23^, and pattern completion ^23^. Additionally, a previous study in mice with 4 copies of the *Gldc* gene encoding the glycine-degrading enzyme glycine decarboxylase, a gene that is expressed mainly in astrocytes, found that these mice have a reduced extracellular glycine level in the DG and display behavioral phenotypes including deficits in working memory, pre-pulse inhibition, startle habituation, social interaction and latent inhibition^24^. Currently, it is unknown to which degree, if any, astrocytes in the DG modulate DG neuronal circuits and behavior mediated by the DG. We hypothesized that manipulation of astrocytic activity in the DG would affect performance in behavioral tasks linked to the neuronal circuitry in the DG, similar to how modulation of astrocytic activity in CA1 affects behavioral phenotypes mediated by neuronal circuits in the CA1 subregion ^10,18^. Given the importance of the DG for hippocampal function, we set out to investigate potential contributions of DG astrocytes to cognitive and emotional behaviors.

## Material and methods

### Animals

All experiments were performed in accordance with the Guide for the Care and Use of Laboratory Animals of the National Research Council and approved by the Institutional Animal Care and Use Committee (IACUC) of the University of Illinois Urbana-Champaign. Our studies used C57BL/6J mice directly purchased from Jackson Laboratory or mice locally bred from mice purchased from the Jackson Laboratory (within 5 generations from the Jackson Laboratory mice). All animals were 3-8 months old at the start of the experiments. Animals were randomly assigned to each experimental group (Fig. 1B). Animals assigned to behavioral experiments were separated into 4 batches, all animals in the same batch have a similar age (within 2 months).

**Fig. 1.**
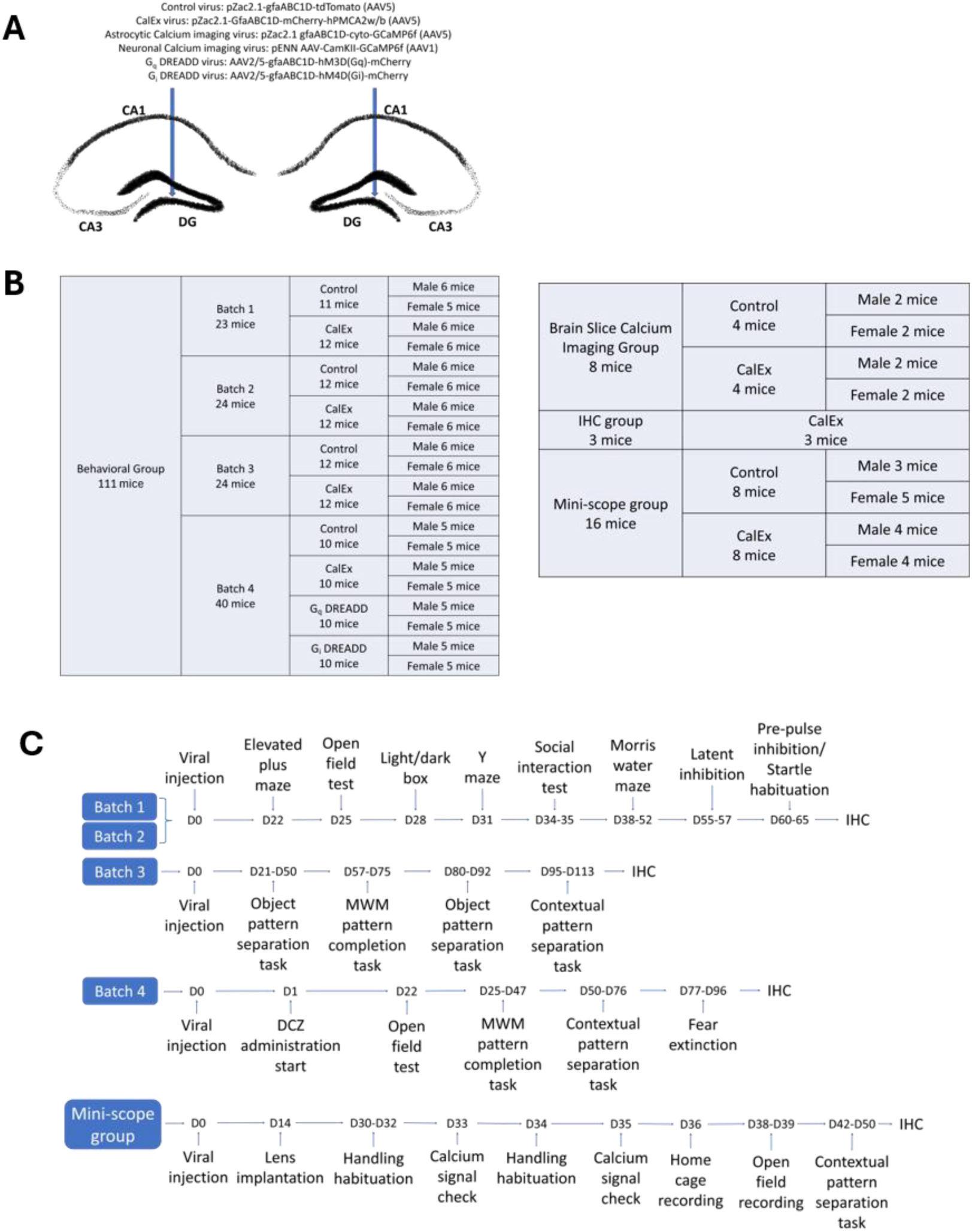
Viral vector injection, experimental group assignments and schedule. (**A**) Schematic of viral vector injection to the dorsal DG. (**B**) Experimental group assignments. (**C**) Schedule of experiments for each behavioral experiment batch.

### Stereotaxic Injection

Mice were deeply anesthetized with isoflurane (2-3% vol/vol) and placed in a stereotaxic frame. Mice were kept warm throughout the procedure with a heating pad. The eyes were covered with eye ointment (equate). A small hole was drilled, and a nano-syringe was inserted bilaterally based on the coordinate relative to bregma (ventral: -2.2mm, lateral: +/-1.4 mm, anterior: -2.1 mm). Viral vector (400nl per side) was injected at 60nl/min and the needle was held at position for 15min after the injection was completed before it was slowly removed. Viral vectors used were pZac2.1-GfaABC1D-tdTomato (#44332-AAV5) purchased from Addgene as control virus, pZac2.1-GfaABC1D-mCherry-hPMCA2w/b (#111568-AAV5) purchased from Addgene as CalEx virus, pZac2.1-GfaABC1D-cyto-GCaMP6f (#52925-AAV5) purchased from Addgene for brain slice calcium imaging, AAV2/5-GfaABC1D-hM3D(G_q_)-mCherry (SKU: construct-1776-aav2-5) purchased from Neurophotonics as G_q_ DREADD virus and AAV2/5-GfaABC1D-hM4D(G_i_)-mCherry (SKU: construct-1777-aav2-5) purchased from Neurophotonics as G_i_ DREADD virus (Fig. 1A), pENN AAV-CamKII-GCaMP6f (AAV1) purchased from Addgene for in vivo calcium imaging. Viral vectors were diluted if needed to maintain a viral concentration close to 1×10^13^vg/mL. After viral injection, mice were kept in single cages for a week before returning to the original group cage except that the mini-scope group were kept single housed to avoid lens damage. Based on previous studies, experiments were performed at least 3 weeks after the viral injection ^18,19^. For animals in batch 1 and 2, the accuracy of viral injection was examined by immunofluorescent to confirm viral expression in the DG after the behavioral experiments. All 47 animals tested showed bilateral expression of viral vector in the DG except 2 control animals showed weak expression on one side.

### DCZ administration

For animals in batch 4, starting the day after viral injection, their home cage water supply was replaced with water-soluble deschloroclozapine dihydrochloride (DCZ) (Hello Bio, Inc., catalog: HB9126) until they were sacrificed. A dose of 3 µg/kg/d was used based on a previous study ^16^. To maintain this dose, the solution was refilled every 2-4 days, consumption volume for the cage and body weight of the animals were recorded at each refill. Concentration of DCZ in the drinking water was adjusted accordingly.

### Behavioral experiments

Behavioral experiments for each batch followed the general timeline outlined (Fig. 1C).

### Elevated plus maze

A plus maze with 30cm arm length was set up 102cm above the floor. The arena was illuminated from above (50 lux measured at the center of the arena). Two opposite arms are walled and the other two are not. Mice were started at the center area and allowed free exploration for 5 minutes. Percentages of time spent in the open arms and the closed arms were analyzed. Animal movement was recorded from above which was tracked and analyzed using the EthoVision XT 15 software (Noldus).

### Open field test

A square (41cm x 41cm x 30cm) white box arena was used as an open field to test for anxiety-like behavior or any potential change to locomotor activity after DCZ administration. The arena was illuminated by a diffuse light source from above (150 lux measured at center). The animals were started at the center of the arena and allowed free exploration for 10 minutes. The central 25% of the arena was designated as “center zone”, the rest as “outer zone”. Animal movement was recorded from above which was tracked and analyzed using the EthoVision XT 15 software (Noldus).

### Light/dark box

The same square white box used in the open field test was used together with a dark insert for light/dark box experiment to test anxiety-like behavior. The dark insert covers approximately 33% of the box and blocks all light for that side. The arena was illuminated from above (100 lux measured at light side). A door at the bottom allows mice to travel freely between the illuminated side of the box and the dark side of the box. The animals were started in the dark side of the arena and allowed free exploration for 10 minutes. Animal movement was recorded from above which was tracked and analyzed using the EthoVision XT 15 software (Noldus).

### Y-maze

A Y-shaped maze (gray, arms: 30 cm length x 8 cm width x 15 cm height) was used to test the spatial memory of the mice. The arena was illuminated from above (100 lux measured at the center of the arena). Each animal was started on the same arm of the Y-maze and allowed free exploration for 8 minutes. Entries into each arm were video recorded and analyzed. Animals with normal spatial memory will remember the arms previously visited and show a tendency to alternate ^25–27^. The percentage of “correct”, i.e., alternating arm entries was calculated for each animal.

### Social interaction test

A transparent three-chamber arena (41cm x 63cm x 21.5cm arena divided into 3 equal chambers, illuminated from above to 100 lux) was used to test the sociability and social preference of mice. Holes are available between each chamber to allow mice free access to all chambers at any time. The arena was placed on a platform surrounded by black curtains to block unintended environmental factors. Each animal was given 5 minutes to explore the arena with an empty wire cup in both the left and the right chamber. Then, a stranger mouse was placed under one wire cup and the experimental mouse was given 10 minutes to explore. Finally, another stranger mouse was placed under the other wire cup, and the experimental mouse was given another 10 minutes to explore. All stranger mice are sex-matched and of similar age as the experimental mice. Animal movement was recorded from above which was tracked and analyzed using the EthoVision XT 15 software (Noldus).

### Morris water maze

A circular plastic water tank 101cm in diameter was used for Morris water maze. The arena was divided into 4 quarters and 4 distinct positional cues were attached to the wall of the water tank between quarters above the water. A platform 10.5cm in diameter was placed in the center of quarter A and submerged 1cm under water. White non-toxic paint was mixed in the water to make the platform invisible. During the initial learning phase, the mice were gently released into the water at the center of one of the quarters that does not have the platform. Mice then were allowed up to 1 minute to reach and stay (more than two seconds) on the hidden platform. If the mice failed to do so within 1 minute, the mice will then be placed on the hidden platform. After reaching and staying on the platform, the mice were left on the platform for 15 seconds before being returned to their cage. This trial was then repeated two times with a different starting quarter for every mouse every day one hour apart until the average latency of reaching and staying on the platform for all experiment groups dropped below 30 seconds for two consecutive days. After the initial learning trials, a probe trial then took place on the next day. The platform was removed during the probe trial and mice were returned to their cage after the 1-minute exploration. The reversal learning phase started one day after the first probe trial. The reversal learning phase followed the same rule as the initial learning phase but with the platform moved to a neighboring quarter. After 6 days of reversal learning, a second probe trial then followed the reversal learning phase. Animal movement was recorded from above which was tracked and analyzed using the EthoVision XT 15 software (Noldus).

### Latent inhibition to conditioned freezing

We measured latent inhibition to conditioned freezing using chambers from Med-Associates, Inc. (Chamber dimension: 26cm x 32cm x 25cm, 200 lux when light switched on). Animals in each group, which is defined by the specific viral vector that was injected, were divided into a preexposure subgroup and a non-preexposure subgroup. On day 1, animals in the preexposure subgroup were presented with pure tones while animals in the non-preexposure subgroup were not. On day 2, 5 times 80 seconds apart, all animals received a pure tone followed by a two second 0.7 mA foot shock. On day 3, all animals were presented with 6 minutes of pure tone after 3 minutes in the chamber. The contextual setups (floor, roof, and odor) of the cages were different for all 3 days to avoid freezing behavior being triggered by anything other than tone. Freezing behavior to the pure tone on day 3 was analyzed for latent inhibition. Freezing behavior to the pure tone was measured as [(freezing time during the tone)/(tone duration) *100%] - [(freezing time before the tone start)/(3 minutes)*100%].

### Pre-pulse inhibition/startle habituation

A previous study revealed that mice with 4 copies of the *Gldc* gene, which is mostly expressed in astrocytes, displayed reduced pre-pulse inhibition and startle habituation ^24^. These mice also displayed reduced extracellular glycine levels and an LTP deficit in DG but not in CA1 ^24^, suggesting that the DG may play a role in modulation of latent inhibition^28^. Furthermore, a previous study has linked the absence of α5-containing GABA_A_ receptors in DG granule cells to latent inhibition deficits ^21^, supporting the idea that the DG modulates latent inhibition. Moreover, reduced expression of a point-mutated GABA_A_ receptor α5 subunit in the hippocampus has also been found to result in pre-pulse inhibition deficits ^29^. A Startle Reflex system (Kinder Scientific, Inc.) (Chamber dimension: 40cm x 35cm x 49cm, not illuminated) was used to evaluate pre-pulse inhibition of acoustic startle following an established protocol ^24^. In short, the animals were habituated to the restrainer, the restraint and the chamber in the first 3 days of the experiment. On day 4, the startle input-output function was measured from 73 dB to 120 dB. On day 5, pre-pulse inhibition and startle habituation data were collected. Average startle responses were measured from 12 120 dB startle pulses scattered throughout the experiment. Percentage pre-pulse inhibition was calculated as the percentage of reduction of response to the startle pulse with 73-85 dB pre-pulses 60ms or 120ms prior to the startle pulse. Startle habituation data were calculated based on 6 startle pulses that were presented at the beginning and 6 startle pulses presented at the end of the experiment.

### Object pattern separation

We used a previously established protocols to test object pattern separation for our animals in batch 3 ^30^. Experimental procedure and data analysis closely followed this established protocol. A gray plastic circular arena 50cm in diameter was used as the arena. The arena was illuminated from above (100 lux measured at the center of the arena). Two sets of identical cylindrical objects approximately 7.5cm in diameter were used as objects. Habituation and training protocol followed the established protocol ^30^. In short, after the animal is trained, the animal is given 5 mins to freely explore and is expected to spend more time investigating the object that is moved depending on the distance moved. Animal investigation of an object is defined as when the animal center point is within 20cm of the object and the head is pointing at the general area of the object (defined as 10cm area from the center of the object). Shift 0 represents no displacement and shifts 1 through 4 represent possible locations with increasingly further displacement. Possible locations for the objects are separated by 4.5cm. Trials with animal spending less than 10s exploring objects are discarded. Animal movement was recorded from above which was tracked and analyzed using the EthoVision XT 15 software (Noldus).

### MWM pattern completion

We used an established protocol based on the Morris water maze to test for pattern completion capability ^23^. After training the animals in the MWM as described above, multiple probe trials were conducted with different numbers of positional cues present. The number of positional cues present followed the order: 4 cues, 1 cue, 2 cues, 4 cues and 0 cue. Only 1 probe trial was performed each day. Animal movement was recorded from above which was tracked and analyzed using the EthoVision XT 15 software (Noldus).

### Contextual pattern separation

To evaluate contextual pattern separation, we used the same chamber used in our latent inhibition experiment following an established protocol used in a previous study that investigated dentate granule cells functions ^23^. In short, after three days of context fear conditioning with foot shock (2 seconds 0.7mA delivered after 3 minutes), the mice were introduced to a different context on day 4 and day 5. After that, mice were trained every day to distinguish the shock context A (with roof insert, acid odor) and the no-shock context B (no roof insert, mint odor). Every two days were analyzed together as a block. For batch 4 animals, in block 10 and in block 11, context B was modified to have the same odor cue as context A (Context B+). Discrimination ratio was calculated as described in a previous study ^21^.

### Fear extinction retention/acquisition

Established studies have already identified that fear extinction acquisition and retention are different learning processes that could be differentially regulated^31^. Thus, our protocol examines both fear extinction acquisition and retention. For batch 4 animals, fear extinction retention/acquisition experiment was conducted 2 days after the contextual pattern separation experiment. The mice spent 4 minutes per day (same as with preceding contextual fear conditioning) every day in the previously shocked context A without receiving any shock. 8 days after day 14, the mice were kept in context A for 30 minutes continuously. This single long session was analyzed in 5 minutes blocks. Animal freezing was recorded and analyzed using chambers from Med-Associates, Inc.

### Brain slice calcium imaging

To evaluate the effect of CalEx virus expression on astrocytic calcium signaling. We injected mice with pZac2.1 gfaABC1D-cyto-GCaMP6f (AAV5) virus in addition to Tdtomato virus or CalEx virus. The ex vivo astrocyte Ca^2+^ imaging with confocal microscopy was performed based on established protocol from published study^32^. Briefly, the animal was decapitated, and the brain was harvested immediately. The brain was sliced into 200µm slices with a vibratome in oxygenated cutting solution (30mM NaCl, 4.5mM KCl, 1.2mM NaH_2_PO_4_, 26mM NaHCO_3_, 10mM D-glucose, 194mM sucrose, 1mM MgCl_2_). The slices were incubated in oxygenated artificial cerebrospinal fluid (ACSF) (124mM NaCl, 4.5mM KCl, 1.2mM NaH_2_PO_4_, 26mM NaHCO_3_, 10mM D-glucose, 2mM CaCl_2_, 1mM MgCl_2_) at 30-32℃ for 30 mins before cooling down to room temperature. Then brain slice was kept in flowing oxygenated ACSF and viewed under a confocal microscope. After locating and focusing on active astrocytes, the imaging session started. First the bathing solution was switched to oxygenated 0.25µM TTX in ACSF for 3 minutes to block neuronal activity, then to oxygenated 10µM phenylephrine (PE) in ACSF for 2 minutes to trigger calcium signaling in astrocytes. Finally, the bathing solution was switched back to oxygenated ACSF and recorded for another 5 minutes. Calcium images were captured at 1 frame/second. ΔF/F was calculated based on the first 10 frames of the recording as baseline. The astrocytes were divided into soma and branches. Both regions were analyzed separately and combined.

### In vivo calcium imaging

For in vivo calcium imaging, the nVista system from inscopix (Bruker) was used. Gradient-index(GRIN) lens implantation was performed two weeks after viral injection. The surgical procedure generally followed published protocol^33^ with some modifications. Primarily, the lens implantation is performed 2 weeks after the virus injection. Calcium signal checks started at least 2 weeks after lens implantation, signal checks continue until clear calcium signal was observed or the animal is excluded due to no signal is observed. After the animals passed signal check, they were used for a recording in home cage for 10mins, 2 sessions of open field recordings (10mins) and contextual pattern separation recordings (identical to protocol previously described) with 1 day of rest between experimental protocols. However, some animals unexpectedly started to develop seizures when the open field recording started. Some mice unexpectedly died later presumably due to server seizures (Fig. 2B). Thus, the contextual pattern separation recording was aborted, and all remaining animals were sacrificed and perfused to preserve brain samples. Detailed schedule for each individual animal is available as part of raw data. To assist analysis for these unexpected seizures issues, 4 extra animals that received lens implantation but were not intended for data collection were also used for IF staining (Fig 2). All 4 animals were injected with GCaMP virus and 3 of them also received CalEx virus.

**Fig. 2.**
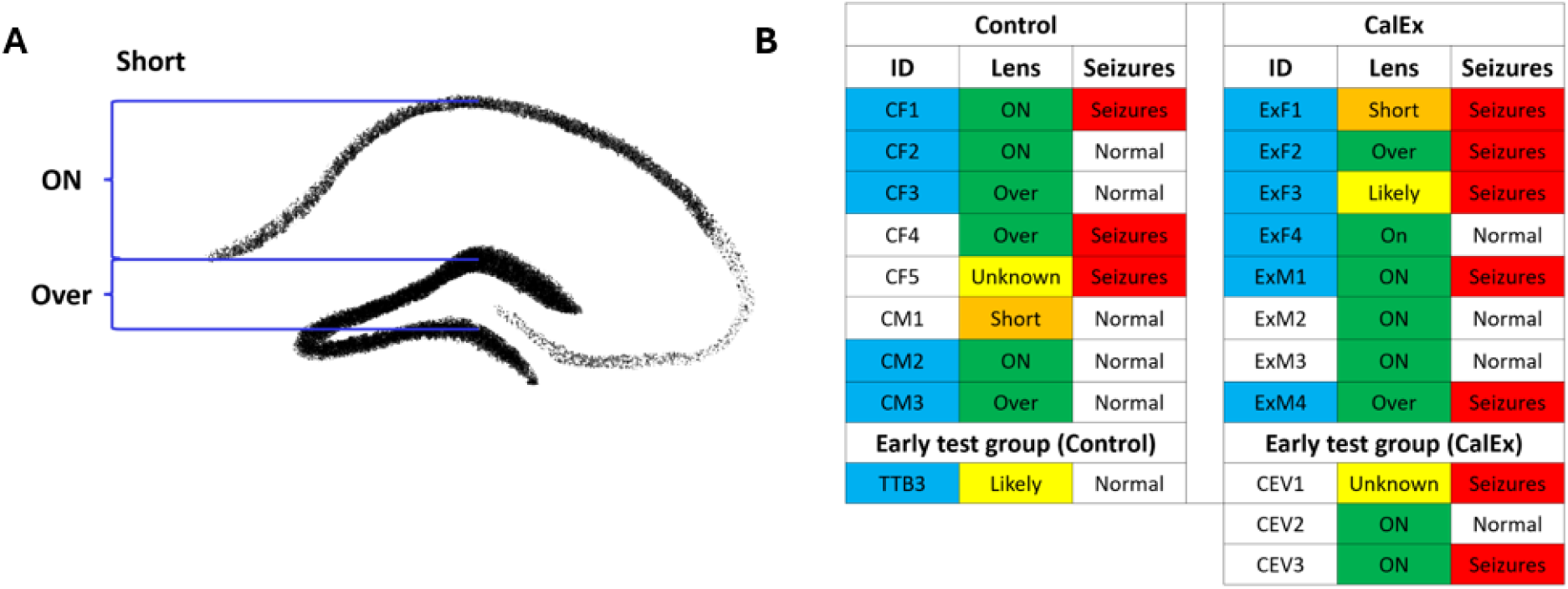
Lens placement and seizure assessment. **(A)** Illustration of lens implantation grading standard. Only “On” or “Over” are considered valid. **(B)** Terminal status for all inspected animals used for in vivo calcium imaging showing animal ID, lens implantation grading, whether the animal shows calcium signal (indicated by blue color) and whether the animal was observed to have seizures.

### Immunohistochemistry (IHC) / Immunofluorescence (IF)

Before staining, animals were put under deep anesthesia with 139mg/kg ketamine mixed with 21mg/kg xylazine and perfused with saline followed by 4% PFA. After submerging the harvested brain in PFA and 30% sucrose for at least 1 day, brains were cut into 40µm slices with a cryostat. The following primary antibodies were used: Rabbit anti-RFP antibody (Rockland #600-401-379), Mouse anti-NeuN antibody (Millpore #MAB377), Mouse anti-S100β antibody (Sigma #2532), Rabbit anti-DCX antibody (Cell Signaling # 4604S), Rabbit anti-ΔFosB (Cell signaling #14695), Rabbit anti-c-fos (Cell Signaling Tech, cat # 2250). Secondary antibodies used are Alexa Flour 568 goat anti-rabbit (Invitrogen A11036), Alexa Fluor Plus 488 goat anti-rabbit (Invitrogen A32731) and Alexa Fluor 488 goat anti-mouse (Invitrogen A11001). DAPI is also used for counterstaining. Cell count is conducted manually with ImageJ after the file name is randomized by program.

For animals used in in vivo calcium imaging, GCaMP expression and lens placement were inspected to determine if the recording from each animal is suitable for analysis (Fig. 2A). Brain samples from some animals were lost due to unexpected animal death. Data from any animal that can not be validated by IHC or was still doubtful after IHC inspection was also analyzed but marked “likely” since almost all viral injections to the DG have been successful.

### Analysis of in vivo calcium imaging

The data from in vivo calcium imaging were analyzed with Inscopix Data Processing Software (IDPS) (Inscopix) after IHC staining confirmed that the lens was implanted at the correct location in the dentate gyrus and that GCaMP was expressed in the dentate gyrus.

### Statistical analysis

ANOVA (one-way or two-way) was used to compare multiple data groups that involve multiple factors followed by Tukey’s multiple comparisons test for most post-hoc analysis except Dunnett’s multiple comparisons test was used when comparing data to one control measurement. Repeated measure variant of ANOVA was used as needed. For one-way ANOVA, Shapiro-Wilk test was used to test data for normality. For repeated measure two-way ANOVA, statistical tests for data normality were not available, thus a QQ plot was visually inspected to check for data normality. If the data violates normality assumption log transformation was applied. Greenhouse-Geisser correction was applied for the repeated measure ANOVA. For comparison between two groups, paired t-test, Welch’s t-test (for unpaired comparisons), Wilcoxon test (for paired data sets that failed the normality test) and Mann-Whitney test (for unpaired data set that failed the normality test) were used. GraphPad Prism 10 software was used for statistical analysis. For z-score analysis, the z-score of each animal in each parameter was calculated as (raw value-average of control)/(standard deviation of control), average z-score was then calculated from all parameters and analyzed with ANOVA.

## Results

### Validation of specificity and effectiveness of CalEx expression in astrocytes in the dentate gyrus

In order to elucidate the function of astrocytes in the DG, we generate mice that express the calcium extrusion pump hPMCA2w/b (CalEx) in these cells. Expression of CalEx results in a disruption of calcium-dependent signaling and thus functional inhibition of the astrocytes. To confirm expression of CalEx in astrocytes in the dentate gyrus after viral transduction with AAV-GfaABC1D-mCherry-hPMCA2w/b, we used immunostaining of CalEx, and of the astrocytic marker S100β, as well as the neuronal marker NeuN. Consistent with expectations, we observed that CalEx expression colocalized with the astrocyte marker S-100β, and not with the neuronal marker NeuN (Fig. 3A), confirming the specificity of the GfaABC1D promoter for astrocytes. To assess the effect of CalEx expression in astrocytes on calcium signaling, brain slice calcium imaging was performed. An astrocytic calcium signal was induced by the α1-adrenergic receptor agonist phenylephrine (PE). This signal was significantly reduced by the expression of CalEx (for soma Mann-Whitney test, p=0.0001, for whole cell Welch’s t-test, p<0.0001, for branch Mann-Whitney test, p<0.0001) (Fig. 3B, C). Overall, these results show that CalEx is expressed specifically in astrocytes and effective at reducing astrocytic calcium signaling.

**Fig. 3.**
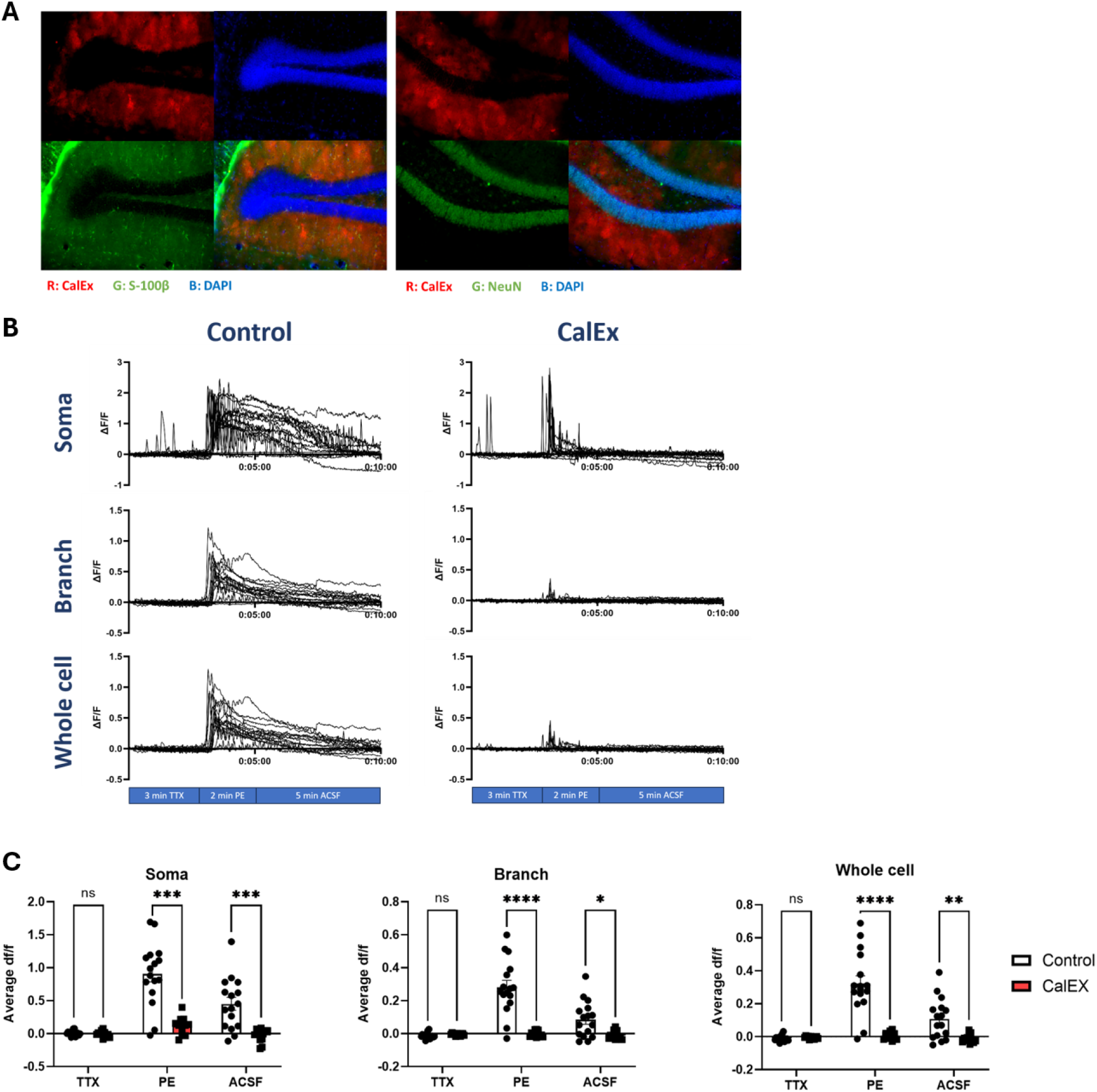
Validation of astrocytic specificity of virus expression and functionality of CalEx expression in the DG. (**A**) Immunofluorescence images showing that CalEx colocalizes with the astrocytic marker S-100β but not with the neuronal marker NeuN. (**B**) Traces of calcium signal from brain slice calcium imaging. (**C**) Average intensity of calcium signal from soma, branch and whole cell at each part of recording. Calcium signal is reduced by CalEx in the entire astrocyte including soma and branches. n=16 cells from 10 brain slices collected from 4 mice for the control group; n=12 cells from 9 brain slices collected from 3 mice for CalEx group. Welch’s t-test (data sets that passed the normality test) or Mann-Whitney test (data sets that failed the normality test), *p < 0.05, **p < 0.01, ***p < 0.001, ****p < 0.0001, Data presented as mean ± SEM.

### Expression of CalEx in dentate gyrus astrocytes marginally improved Morris Water Maze (MWM) performance

A previous study has shown that knockout of inhibitory α5-containing GABA_A_ receptors in DG granule cells impairs reversal learning in the MWM^21^, suggesting the reversal learning is modulated by these neurons of the dentate gyrus. Thus, we used the MWM to examine whether expression of CalEx in DG astrocytes would affect reversal learning in the MWM. Somewhat surprisingly, the CalEx mice showed a significantly shorter latency to reach the platform on one day during both the initial training phase and the reversal learning phase (Fig. 4) (Tukey’s multiple comparisons test, for day 4 during training phase p=0.0015, for day 3 during reversal phase p=0.0404). In both phases the average latency to reach the platform for CalEx mice was lower than that of the control mice for most days. However, there was no statistically significant difference between CalEx mice and control mice for both phases (for training phase Mixed-effect model, for factor virus, p=0.0947, F=2.915, for factor day p<0.0001, F=14.99, for interaction between factors p=0.1369, F=1.662, for reversal phase Two-way ANOVA for repeated measures, for factor virus, p=0.1541, F=2.101, for factor day p<0.0001, F=6.857, for interaction between factors p=0.1795, F=1.593). Thus, our results indicate that chronic inhibition of DG astrocytic calcium activity leads to a small and temporary improvement in spatial learning and reversal learning.

**Fig. 4.**
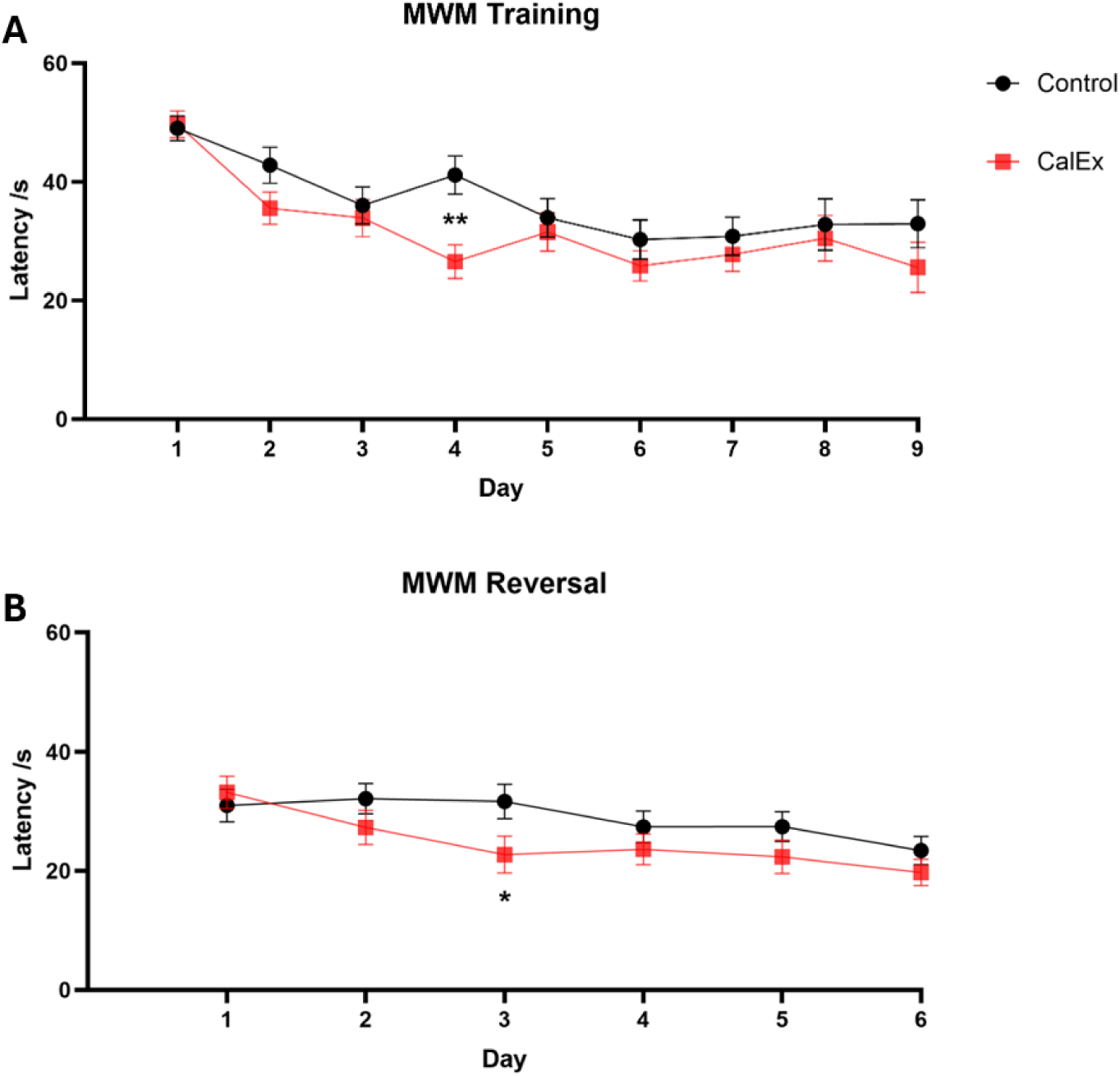
CalEx expressed in DG astrocytes improves initial learning and reversal learning in the MWM. (**A**) Latency to reach the platform during the initial learning phase. CalEx mice showed a significantly shorter latency on day 4 of training. (**B**) Latency to reach the platform during the reversal learning phase. CalEx mice showed significantly shorter latency compared to the control on day 3 of training. For (**A**) Mixed-effect model, n=23-24 for day 1-7, n=12 for day 8-9, For (**B**) Two-way ANOVA repeated measures, n=23-24, *p < 0.05, **p < 0.01. Data shown in this figure originate from behavioral group batch 1 and batch 2. Data presented as mean ± SEM.

### CalEx mice displayed improved pattern separation

Many studies have shown that the DG is critical for pattern separation ^21,23,34,35^. We were interested in the question whether astrocytes in the dentate gyrus contribute to pattern separation, thus we used both an object pattern separation protocol and a contextual pattern separation protocol to investigate whether expression of CalEx in DG astrocytes would affect the capability to perform object and contextual pattern separation^23^. For object pattern separation, the CalEx mice showed a significantly improved discrimination index starting with shift distance 1 (Mixed effect model, p<0.0001, F=30.1, Dunnett’s multiple comparisons test, p=0.038 at shift distance 1), compared to control mice which started to show a significant improvement at shift distance 2 (Fig.5 B-C) (One way ANOVA repeated measures, p<0.0001, F=53.4, Dunnett’s multiple comparisons test, p=0.3089 at shift distance 1). When the discrimination index at each shift distance is compared between control and CalEx mice, the CalEx mice showed a trend towards an improved performance compared to control mice at shift distance 3 (Tukey’s multiple comparisons test, p=0.0573) (Fig. S1A). Also, the CalEx mice showed a trend towards an increased investigation time difference between the shifted object and the stationary object compared to control mice at shift distance 3 (Tukey’s multiple comparisons test, p=0.0603) (Fig. S1B), combined analysis of both parameters (discrimination index, investigation time) with z-score shows CalEx mice have improved performance at shift distance 3 (repeated measure two-way ANOVA, for factor virus p=0.8831, F=0.02212, for factor shift distance p=0.1133, F=2.023, for interaction p=0.1088, F=2.055, Tukey’s multiple comparisons test, p=0.0397 at shift distance 3) (Fig. S1C). Overall, these findings indicate that CalEx mice have an improved object pattern separation performance compared to control mice. In the contextual pattern separation paradigm (Batch 3, Fig. 1C), the control mice did not show significantly different freezing percentages between contexts until block 2 (Fig. 5E) (for factor context p=0.0009, F=20.19, for factor block p=0.0098, F=4.098, for interaction p=0.0529, F=2.974, p=0.1176 at block 1, Tukey’s multiple comparisons test, p=0.028 at block 2), thus showing contextual pattern separation only starting at block 2. In contrast, CalEx mice displayed rapid pattern separation starting in the first block (Fig. 5F) (for factor context p=0.0003, F=26.39, for factor block p=0.0744, F=2.898, for interaction p=0.0185, F=3.085, Tukey’s multiple comparisons test, p=0.005 at block 1). Thus, inactivation of DG astrocytes by calcium extrusion improves rapid pattern separation. In a separate experiment using animals with a different experimental history (Batch 4, Fig. 1C), control mice showed contextual pattern separation already in block 1 (for factor context p<0.0001, F=51.4, for factor block p=0.0164, F=3.583, for interaction p=0.0729, F=2.427, Tukey’s multiple comparisons test, p=0.0037 at block 1) (Fig. S1D), indicating that the two contexts in this experiment were too distinct for this batch of animals. In these animals, we assessed learned pattern separation that may take several days to develop^36^. Indeed, we observed that while in control mice the discrimination ratios did not change over the duration of the 11 blocks (for factor virus p=0.6221, F=0.5954, for factor block p<0.0001, F=18.28, for interaction p=0.1067, F=1.507, Dunnett’s multiple comparisons test, at block 11 p=0.2844), in the CalEx mice we observed significantly higher discrimination ratios in blocks 11 compared to block 1 (Dunnett’s multiple comparisons test, p=0.0474) (Fig. S2). In this experiment, we also included groups expressing a G_i_ DREADD and a G_q_ DREADD in DG astrocytes, in which G_i_-dependent pathways and G_q_-dependent pathways, respectively, are activated in astrocytes by chronically administering DCZ. G_i_ DREADD mice and G_q_ DREADD mice displayed a significant difference in freezing already in blocks 1 and 2, similar to control mice (for G_i_ DREADD factor context p=0.0016, F=19.82, for factor block p=0.0022, F=6.54, for interaction p=0.0836, F=2.512, Tukey’s multiple comparisons test, p=0.0025 at block 1, for G_q_ DREADD factor context p=0.0001, F=41.9, for factor block p<0.0001, F=12.88, for interaction p=0.002, F=5.358, Tukey’s multiple comparisons test, p=0.0167 at block 1) (Fig. S1D). While in the G_i_ DREADD mice the discrimination ratio did not increase over the course of 11 blocks, in the G_q_ DREADD mice it was increased in blocks 8-10 compared to block 1 (Dunnett’s multiple comparisons test, at block 8 p=0.012, at block 9 p=0.0097, at block 10 p=0.013) (Fig. S2). Overall, our results on learned pattern separation show an increased discrimination ratio across time and thus learned pattern separation in CalEx mice and in G_q_ DREADD mice, but not in control or G_i_ DREADD mice. Astrocytes in the DG thus appear to suppress rapid pattern separation and support learned pattern separation via G_q_-dependent pathways.

**Fig. 5.**
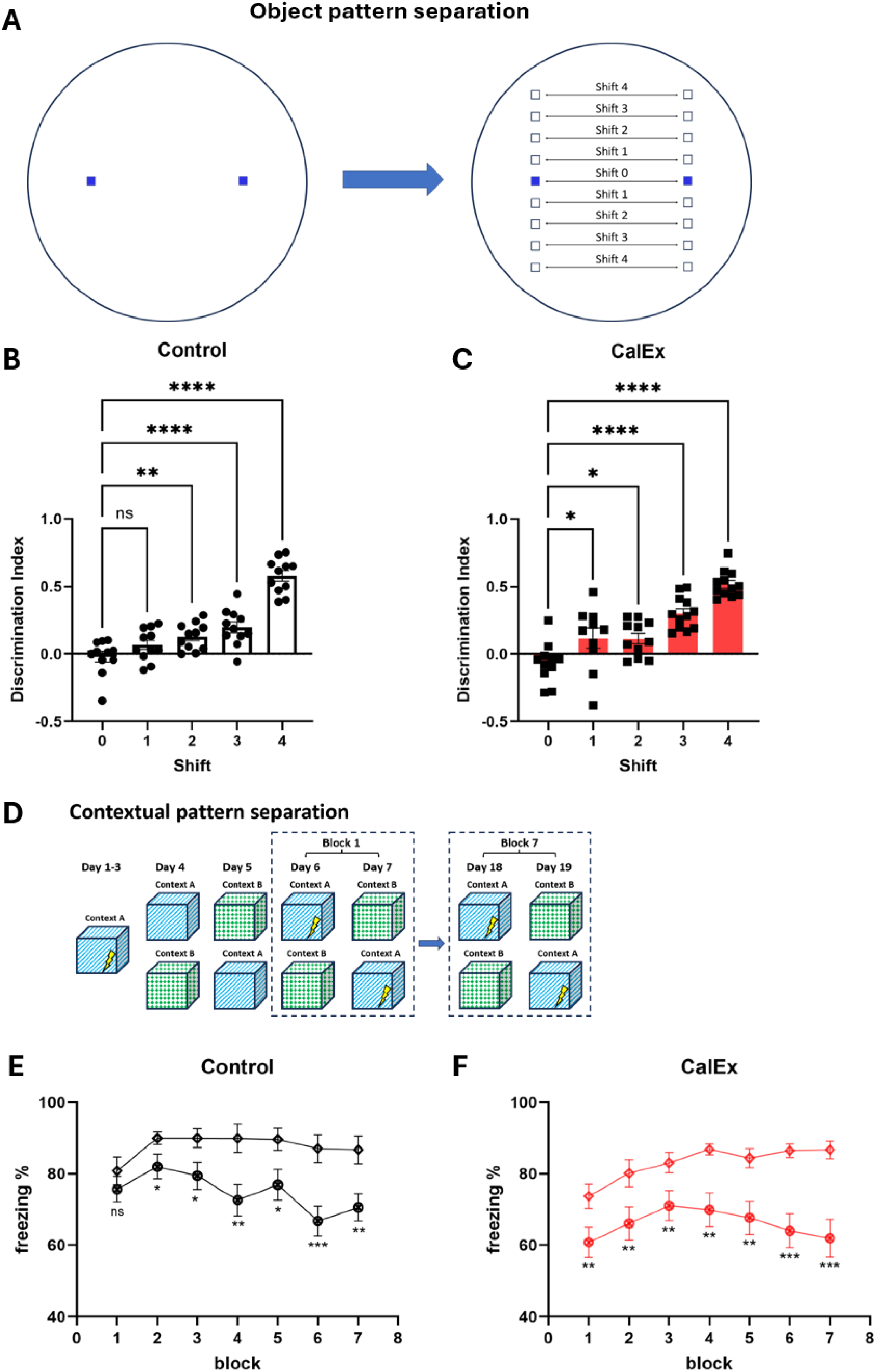
CalEx improves object and contextual pattern separation. **(A)** Illustration of object pattern separation arena setup (**B-C**) Discrimination index from object pattern separation protocol at each object shift distance for control and CalEx mice. (**D**) Illustration of contextual pattern separation protocol. (**E-F**) Freezing percentage in each of the two contexts from contextual pattern separation protocol for control and CalEx mice. One way ANOVA repeated measures was used for (**B**), Mixed effect model used for (**C**). Two-way ANOVA repeated measures was used for (**E**, **F**), n=11 for (**A**), n=10-12 for (**B**). n=12 for (**E**, **F**), *p < 0.05, **p < 0.01, ***p < 0.001, ****p < 0.0001. Data presented as mean ± SEM.

### CalEx mice displayed a deficit in a pattern completion task

Pattern completion and pattern separation have been proposed to be distinct but complementary processes, with improvement in one leading to the impairment of the other ^23,34^. As we have observed improved pattern separation in CalEx mice, we used the MWM to examine the pattern completion capability of the CalEx mice. When each trial condition (i.e., 0, 1, 2, or 4 cues; Fig, 6A) was examined individually, CalEx mice displayed a significant reduction in the number of phantom platform crossings compared to control mice when 2 cues were present (repeated measure two-way ANOVA, for factor cue p=0.0467, F=2.536, for factor virus p=0.1071, F=2.711, for interaction p=0.4145, F=0.9828, Tukey’s multiple comparisons test, p=0.0313) and a trend of shorter phantom platform duration when 2 cues were present (repeated measure two-way ANOVA, for factor cue p=0.0715, F=2.299, for factor virus p=0.214, F=1.592, for interaction p=0.2779, F=1.293, Tukey’s multiple comparisons test, p=0.0784) (Fig. S3E). When we averaged the data from each mouse in both trials with 4 cues into a “full cue” trial condition, and 1-cue and 2-cues trials into a “partial cue” trial condition, we observed that when only partial cues are present CalEx mice displayed a significantly lower number of phantom platform crossings (for factor cue p=0.0024, F=7.086, for factor virus p=0.1471, F=2.182, for interaction p=0.1587, F=1.924, at partial cue p=0.019) and had less time on phantom platform (for factor cue p=0.001, F=7.944, for factor virus p=0.2728, F=1.235, for interaction p=0.0933, F=2.489, at partial cue p=0.0339) (Fig. 6B) if both batches were combined. In contrast, no difference was seen in the full cue condition or in the no cue condition (Fig 6B) (for crossing at full cue p=0.1425, at no cue p=0.9791, for duration at full cue p=0.4552, at no cue p=0.8831). Overall, the results indicate that CalEx mice have a pattern completion deficit compared to controls.

**Fig. 6.**
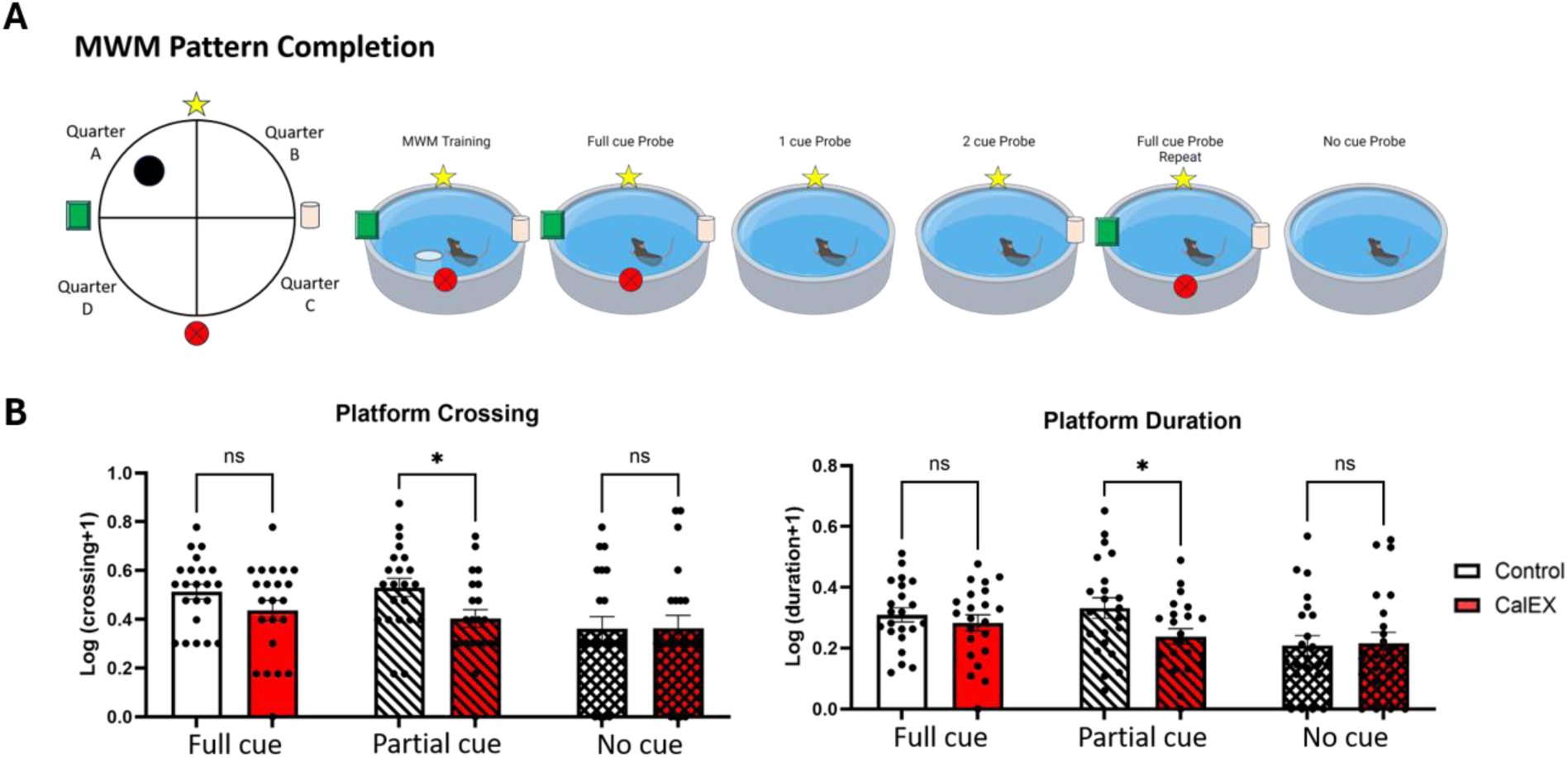
Expression of CalEx in DG astrocytes impairs pattern completion. **(A)** Illustration of MWM setup and pattern completion experiment based on the MWM task. **(B)** Total time spent on phantom platform during probe trial and total number of phantom platform crossings during probe trial with batches 3 and 4 combined, log transformation is used. Two-way ANOVA repeated measures, n=22, *p < 0.05. Data presented as mean ± SEM.

### Chronic G_i_ DREADD and G_q_ DREADD activation differentially induced between-session fear extinction and within-session fear extinction deficits

As fear extinction is also modulated by the DG ^21,22^, after completion of the pattern separation experiment we assessed fear extinction in behavioral group batch 4 (Fig.7A). When comparing the freezing percentages on each of the days 2-14 to those on day 1, only the G_i_ DREADD mice did not show any significant reduction in freezing percentages over the course of 14 days (repeated measure two-way ANOVA, for factor day p<0.0001, F=19.32, for factor virus p=0.1043, for interaction p=0.0587, F=1.536, Dunnett’s multiple comparisons test, for G_i_ DREADD mice on day 14 p=0.087) (Fig. 7B). Although G_q_ DREADD mice showed a significant reduction in freezing on day 3 (Dunnett’s multiple comparisons test, p=0.038), the difference was not significant for the following days until day 7 (Dunnett’s multiple comparisons test, p=0.0124), similar to control and CalEx mice, these mice showed significant reduction in freezing percentage on day 7 or 8 (Dunnett’s multiple comparisons test, for control mice on day 7 p=0.0099, for CalEx on day 8 p=0.0185) (Fig. 7B). It is noteworthy that all experimental groups appeared to have a sudden increase in freezing percentage on day 9. The bedding in the home cages of all animals was replaced after the end of the experiment on day 8. It is conceivable that this disruption, which potentially introduces stress, may have resulted in this brief increase in freezing percentages. Overall, these data demonstrate that while the percentage freezing is reduced on days 7 and/or 8 compared to day 1 for control, CalEx, and G_q_ DREADD mice, G_i_ DREADD mice do not show any decrease in the freezing percentage on days 2-14 compared to day 1 in this testing protocol which is limited to 4 minutes of fear extinction acquisition per day. This finding suggests a between-session fear extinction deficit in the G_i_ DREADD mice. However, when the animals were subsequently subjected to a single 30-minute fear extinction session, the freezing percentages of the G_i_ DREADD mice were reduced within 15 minutes (repeated measure two-way ANOVA, for factor time p<0.0001, F=18.84, for factor virus p=0.255, F=1.413, for interaction p=0.5344, F=0.9175, Dunnett’s multiple comparisons test, at minute 15 p=0.0452), similar to other experimental groups (Dunnett’s multiple comparisons test, for control mice at minute 15 p=0.027, for CalEx mice at minute 15 p=0.0129) except the G_q_ DREADD mice which did not show any significant reduction in freezing percentage (Dunnett’s multiple comparisons test, at minute 15 p=0.0566) (Fig. 7C). After normalizing the freezing percentage to the initial freezing percentages for individual mice in the first 5-min period, control mice, CalEx mice, and G_i_ DREADD mice displayed reduced freezing in more than one of the following 5-min periods (repeated measure two-way ANOVA, for factor time p<0.0001, F=10.03, for factor virus p=0.5639, F=0.6903, for interaction p=0.4757, F=0.9696, Dunnett’s multiple comparisons test, for control mice at minute 15 p=0.0273, for CalEx mice at minute 15 p=0.0317, for G_i_ DREADD mice at minute 15 p=0.0399), while the G_q_ DREADD mice did not display any significant changes in freezing across all 5-min periods for 30 min (Dunnett’s multiple comparisons test, at minute 15 p=0.069), suggesting a deficit in within-session fear extinction in a single 30-min long extinction session (Fig. 7D). Overall, our data suggests that chronic G_i_ DREADD activation in DG astrocytes inhibits between-session fear extinction over a 14-day period with repeated short 4-min sessions, while chronic G_q_ DREADD activation inhibits within-session fear extinction in a single long 30-min session, indicating that these responses are differentially regulated by distinct astrocytic pathways.

**Fig. 7.**
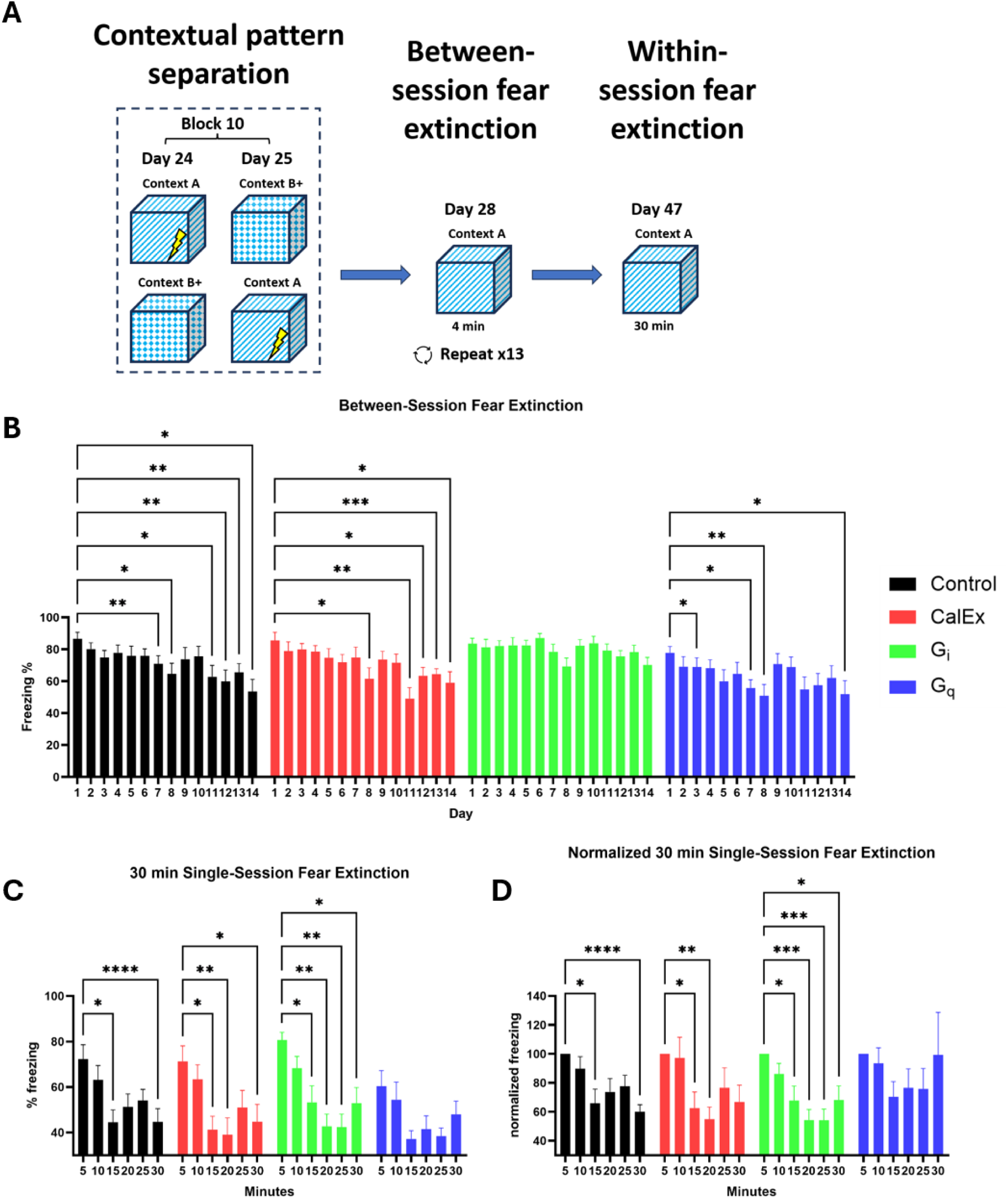
Effect of CalEx, chronic G_i_ DREADD activation and chronic G_q_ DREADD activation in DG astrocytes on between- and within-session fear extinction. **(A)** Illustration of between- and within-session fear extinction protocol after the end of pattern separation experiment. (**B**) Freezing percentage during the repeated short session fear extinction training comparing freezing percentage to the initial freezing on day 1. (**C**) Freezing percentage during the 30-minute single session fear extinction. The graph shows the freezing percentage for each 5-min interval. (**D**) Freezing percentage (same data as in C) normalized to the first 5 minutes of the 30-minute single session fear extinction. Data shown in this figure originate from behavioral group batch 4. Two-way ANOVA repeated measures was used for all panels, n=10, *p < 0.05, **p < 0.01, ***p < 0.001. Data presented as mean ± SEM.

### Multiple behavioral phenotypes are not significantly changed by expression of CalEx or chronic activation of G_i_ DREADD and G_q_ DREADD

Other than pattern separation, pattern completion and fear extinction, DG is also involved in many other behavioral tasks such as locomotion, anxiety-like behavior, social interaction behavior and startle habituation which could be modulated by astrocytes. Thus, we investigated whether our astrocytic manipulation would have an effect in multiple other behavioral tasks. In the elevated plus maze, the CalEx mice overall (male and female combined) did not show a significant difference in the time spent in the open arms compared to control (Welch’s t-test, p=0.5759), but the female subpopulation spent a significantly higher percentage of time in the open arm (Welch’s t-test, p=0.0353), while there was no difference in males (Welch’s t-test, p=0.2928) (Fig. 8A). In the open field test, time in the center was not different between CalEx mice, control mice, G_i_ DREADD mice and G_q_ DREADD mice (for batch 3 Welch’s t-test, for male p=0.167, for female p=0.451, for combined p=0.4638, for batch 4 One-way ANOVA, p=0.8203, F=0.3068) (Fig. 8B, C). Travel distance and average speed in the open field also showed no difference between all experimental groups indicating no locomotion changes induced by expression of CalEx, chronic G_q_ or G_i_ activation in DG astrocytes (for batch 3 Welch’s t-test, for speed p=0.9505, for distance p=0. 9431, for batch 4 One-way ANOVA, for speed p=0.3529, F=1.122, for distance p=0.3571, F=1.112) (Fig. 8B, C). Likewise, in the light/dark box, there was no difference between CalEx mice and control mice in the time spent in the lit area for both sexes analyzed separately and combined (Welch’s t-test, for male p=0.9415, for female p=0.3788, for combined p=0.4999). Thus, while CalEx females may display an reduced anxiety-like phenotype in the elevated plus maze, it was not consistently observed in all tests assessing anxiety-like behaviors. We did not observe any significant difference between the CalEx mice and control mice in the alternation rate in the Y-maze (Welch’s t-test, for male p=0.9711, for female p=0.1058, for combined p=0.2485), in sociability (repeated measure two-way ANOVA, for factor side of chamber p<0.0001, F=130.5, for factor virus p=0.9977, F=8.328×10^-6^, for interaction p=0.6043, F=0.4275, Tukey’s multiple comparisons test, for control mice between stranger and empty cup p<0.0001, for CalEx mice between stranger and empty cup p=0.001) and social preference testing (repeated measure two-way ANOVA, for factor side of chamber p<0.0001, F=56.49, for factor virus p=0.7659, F=0.08971, for interaction p=0.4075, F=0.8882, Tukey’s multiple comparisons test, for control mice between stranger and familiar mice p=0.1487, for CalEx mice between stranger and familiar mice p=0.5056) in the three-chamber social interaction test, in latent inhibition to conditioned freezing (Welch’s t-test, for control mice p=0.0283, for CalEx mice p=0.0066), in startle habituation (paired t-test, for control male p=0.3505, for control female p=0.5792, for control combined p=0.2636, for CalEx male p=0.8492, for CalEx female p=0.6302, Wilcoxon test, for CalEx combined p=0.6033) and in pre-pulse inhibition (repeated measure two-way ANOVA, for factor virus p=0.2451, F=1.387, for factor pre-pulse setting p<0.0001, F=130.6, for interaction p=0.7821, F=0.3574) (Fig. 8E-I). These findings suggest that working memory, social interaction, ignoring irrelevant stimuli, and non-associate learning are normal in CalEx mice. Overall, our data show that our manipulation of astrocytic activity in dentate gyrus induced very selective cognitive deficits as shown above and does not affect other behaviors which have previously been linked to DG function.

**Fig. 8.**
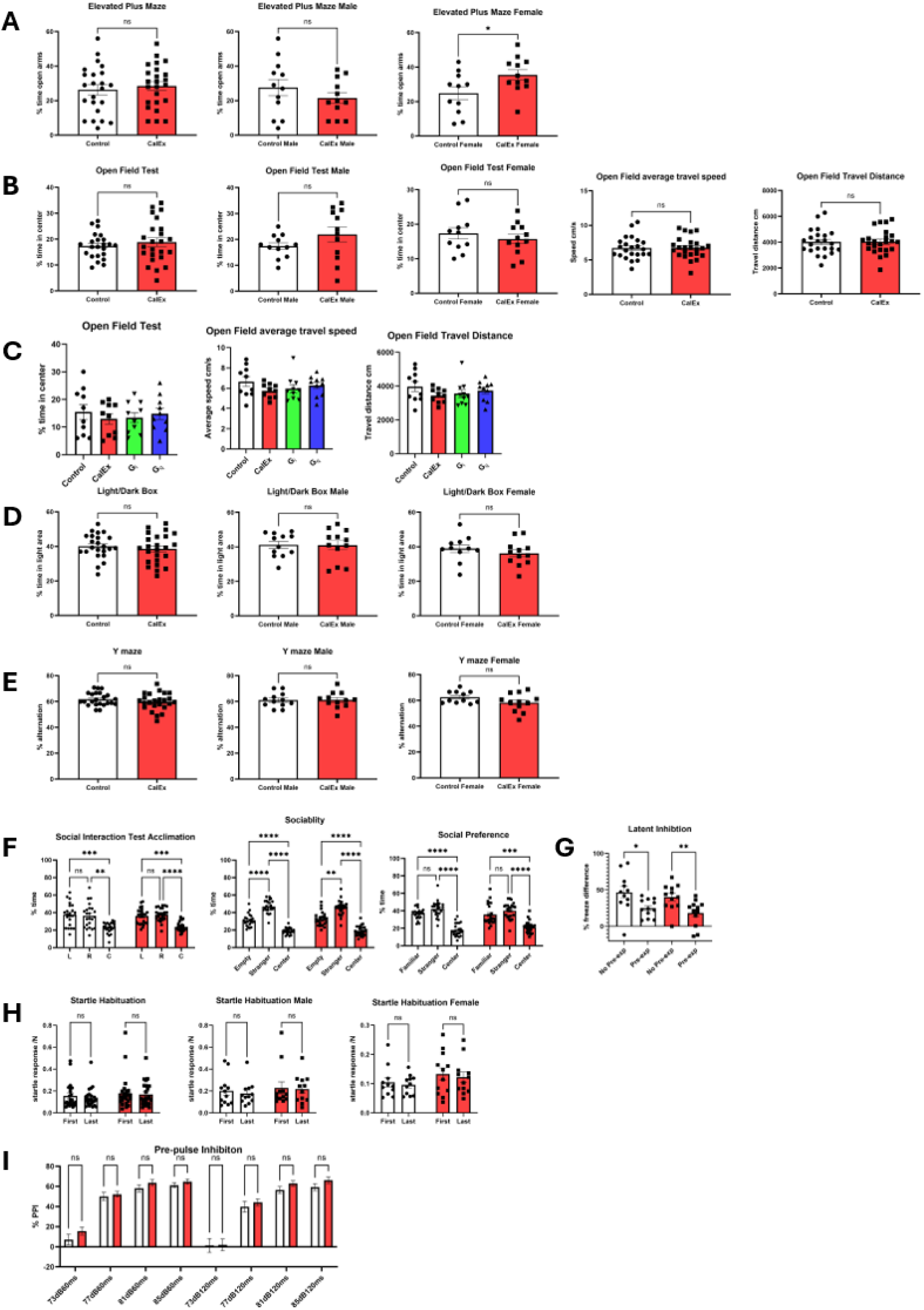
No effect of DG astrocyte manipulation across multiple behavioral domains. Behavioral performance comparison between control, CalEx, G_i_ DREADD and G_q_ DREADD mice in elevated plus maze (**A**), open field (**B, C**), light/dark box (**D**), Y-maze (**E**), three-chamber social interaction test (**F**), latent inhibition of conditioned freezing (**G**), startle habituation (**H**), and pre-pulse inhibition **(I)**. Welch’s t-test for (**A**, **B**, **D**, **E**, **G**), One-way ANOVA for (**C**), Two-way ANOVA for (**F**, **I)**, Paired t-test for (**H**). Data from all panels originate from batch 1 and batch 2 except (**C**) where data originate from batch 4. *p < 0.05, **p < 0.01, ***p < 0.001. Data presented as mean ± SEM.

### CalEx does not invoke significant changes in c-Fos and ΔFosB expression or adult neurogenesis

Since our findings have shown that astrocytic expression of CalEx in mouse dentate gyrus has identifiable effects on cognition, we examined whether expression of CalEx alters the general activity level of neurons in the DG by analyzing expression and density of the activity markers c-Fos, which is an immediate early gene, and ΔFosB, a splice variant of the FosB gene which is considered to indicate a sustained molecular switch due to its long half-life. Also, since dentate gyrus is a subregion known to harbor adult neurogenesis in CNS, we also stained for doublecortin (DCX), a marker for immature neurons, to screen for any potential impact of CalEx expression on adult neurogenesis. For all markers mentioned, both the number of positive cells and the density of positive cells showed no significant difference between CalEx and control groups (for c-fos number Mann-Whitney test, p=0.6159, for c-fos density Welch’s t-test, p=0.764, for ΔFosB number Welch’s t-test, p=0.0992, for ΔFosB density Mann-Whitney test, p=0.2829, for DCX number Welch’s t-test, p=0.8953, for DCX density Welch’s t-test, p=0.7772) (Fig. 9A-C). This result shows that CalEx expression in DG astrocytes does not result in a change of general neuronal activity to a level that would be reflected by c-Fos and ΔFosB, and that adult neurogenesis is not significantly affected. However, this does not rule out potential more subtle changes in neuronal signals that would still be sufficient to induce behavioral changes. To investigate whether such changes may be present in the CalEx mice, we employed in vivo calcium imaging techniques.

**Fig. 9.**
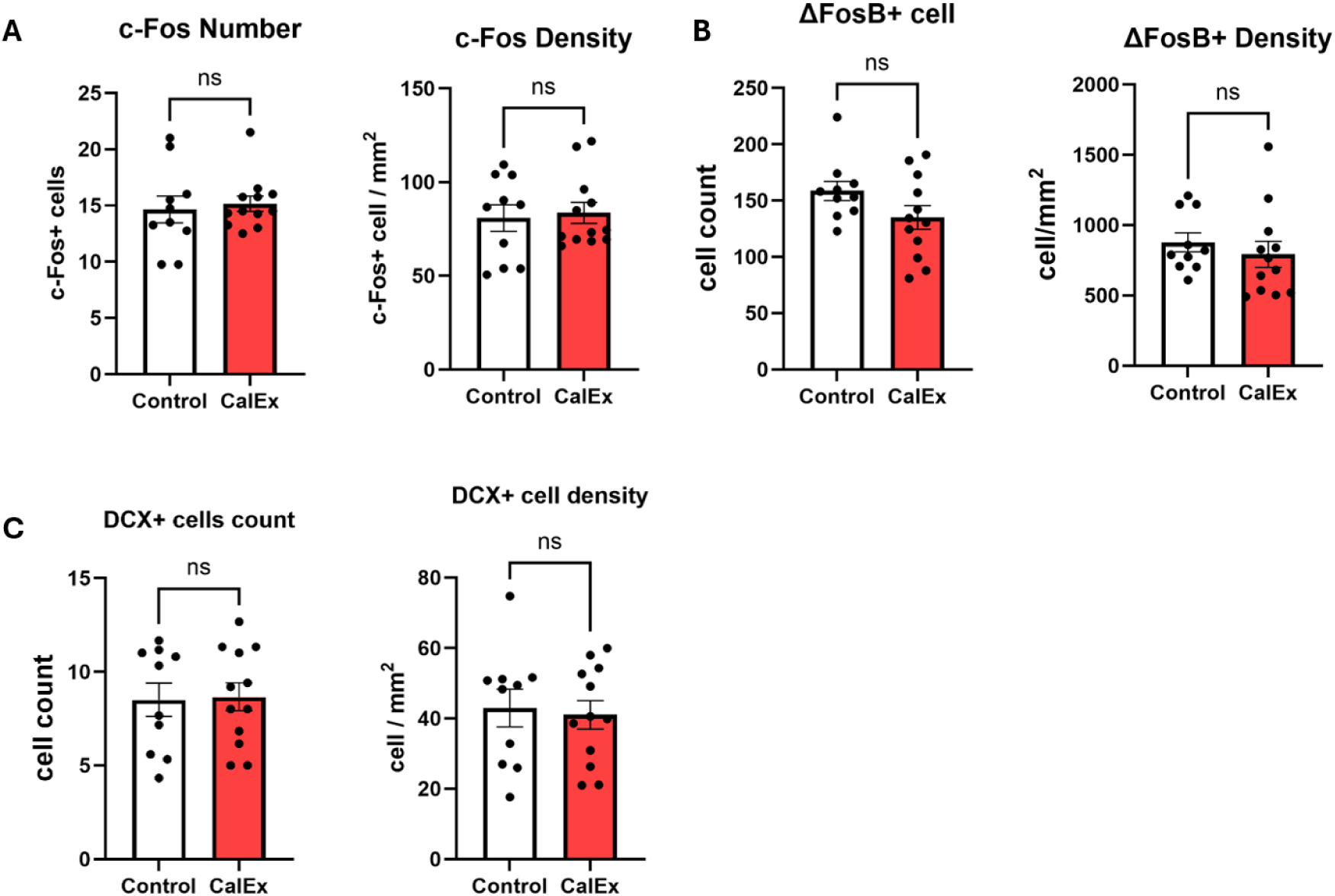
c-Fos, ΔFosB, and doublecortin expression in mice expressing CalEx in DG astrocytes. The number and density of dentate gyrus cells positive for this marker are shown. **(A)** c-fos. **(B)** ΔFosB. **(C)** doublecortin. Welch’s t-test (data sets that passed the normality test) or Mann-Whitney test (data sets that failed the normality test), n=12, all brain samples originate from behavioral batch 3. Data presented as mean ± SEM.

### Expression of CalEx in dentate gyrus astrocytes altered local neuronal calcium signal pattern

We recorded the activity from DG granule cells in behaving animals that had been injected with AAV-CaMKII-GCaMP6f and into which a GRIN (gradient index) lens has been implanted. When calcium activity was recorded in the home cage, the CalEx mice displayed a significantly higher calcium event rate compared to the control mice (Mann-Whitney test, p<0.0001) (Fig. 10A). While the amplitude showed a trend of being increased by CalEx (Mann-Whitney test, p=0.0578) (Fig. 10B), the trend is not observed if the animal with lens implantation marked as “likely” is included (Mann-Whitney test, p=0.3233) (Fig. S4B). Surprisingly, when we looked at the overall average calcium signal intensity level, the CalEx mice had a significantly lower value than the control mice despite a higher calcium event rate (Mann-Whitney test, p=0.0405) (Fig. 10C). This is consistent with the observation that while calcium signals from the control mice tend to have a lower number of events, each calcium event tends to last longer while CalEx mice tend to have a higher number of events, but each tends to form a narrow peak (Fig. 10D). In any case, our in vivo calcium imaging results show that the expression of CalEx in DG astrocytes modulates the calcium signals of dentate gyrus neurons. Opposite changes in calcium event rate (increased) and calcium signal intensity (decreased) suggests that inhibition of the DG astrocyte calcium signals has an effect on DG neuronal signal pattern, which may underlie the observed behavioral phenotype in the CalEx mice.

**Fig. 10.**
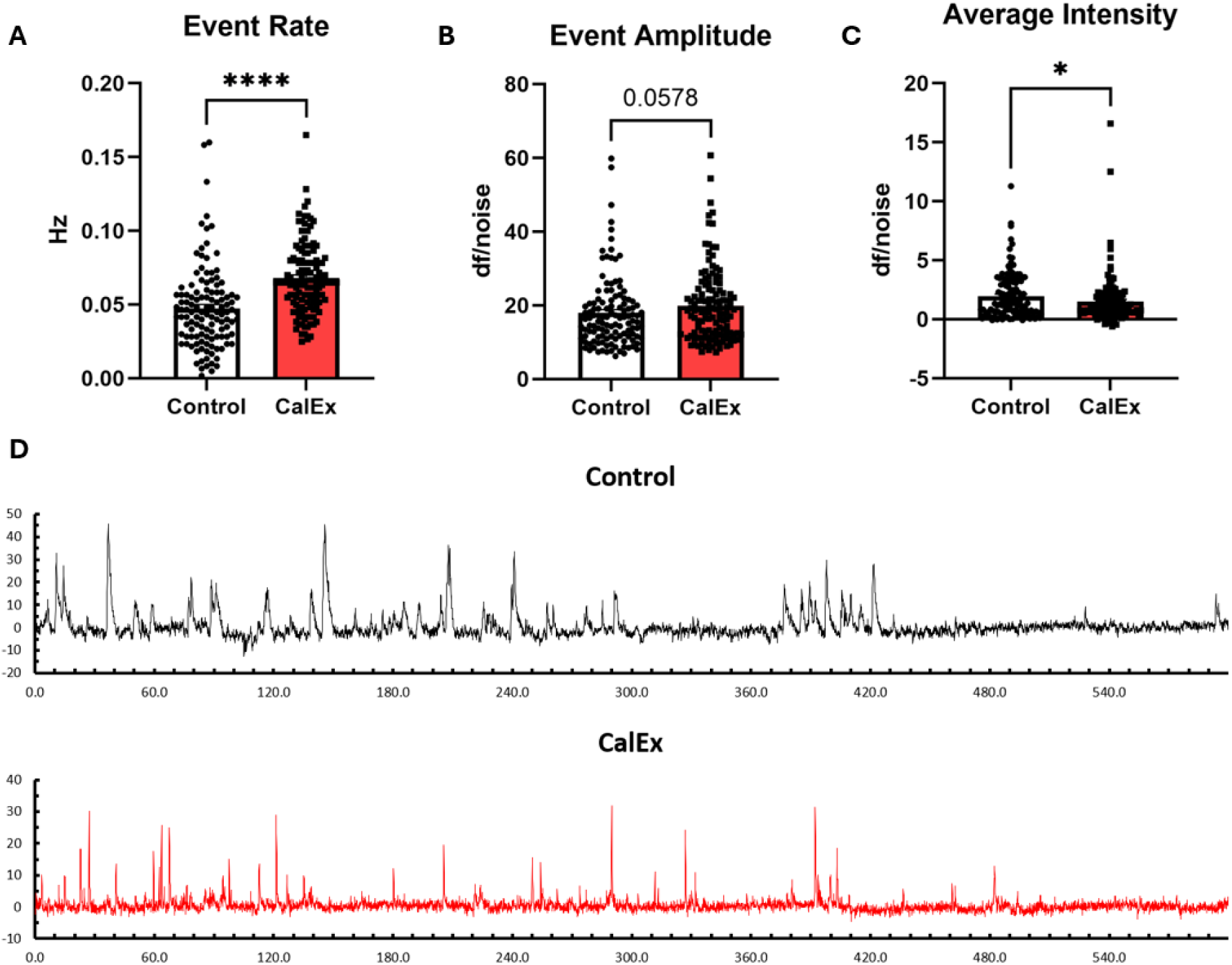
CalEx mice show altered neuronal calcium signal in DG neurons. Comparison of neuronal calcium event rate **(A)**, event amplitude **(B)** and average signal intensity **(C)** between control and CalEx mice. **(D)** Representative sample trace of calcium signaling from control and CalEx animals. Mann-Whitney test, n=121-128. *p < 0.05, ****p < 0.0001. Data presented as mean ± SEM.

## Discussion

Our understanding of diverse regulatory functions of astrocytes in different neuronal networks has advanced rapidly in recent years, to a large part due to the availability of a variety of novel genetic methods allowing us to bidirectionally manipulate the activity of astrocytes^6^. These tools allowed us to establish a basic understanding of the behavioral and neuronal consequences of chronic manipulation of astrocytic activity in the DG. The DG is known to have a significant role in pattern separation and pattern completion ^21,23,34^ and is involved in MWM reversal learning and fear extinction ^21,22,37^, but the contribution of astrocytes to DG function is largely unknown. In the current study, we have shown that inhibition of calcium signaling in DG astrocytes improves pattern separation (Fig. 5) and impairs pattern completion (Fig. 6). It also marginally improves initial learning and reversal learning in the MWM (Fig. 4). Using tools that activate astrocytes, we found that chronic G_i_ DREADD activation impairs between-session fear extinction while chronic G_q_ DREADD activation improves pattern separation learning and impairs within-session fear extinction (Fig. S2, 7). A variety of DG-dependent behaviors were not affected by the modulation of the activity of astrocytes in our study (Fig. 8), underscoring the specificity of the observed responses. Our data from immunofluorescence and in vivo calcium imaging provide a basic understanding of the neuronal mechanism underlying such behavioral changes. While expression of markers of general activity (c-fos or ΔFosB) and of neurogenesis (doublecortin) were unaltered, in vivo calcium imaging revealed that chronic inhibition of DG astrocytes via calcium extrusion alters the calcium activity pattern in DG neurons, with an increased event rate but a reduced average signal intensity, providing a cellular basis for the behavioral changes observed.

Rapid pattern separation deficits have been described previously in mice lacking the GluN1 (NR1) subunit and thus functional NMDA receptors in DG granule cells^35^. In a contextual pattern separation paradigm similar to the one used by us, these mice displayed a lower discrimination ratio starting on day 2 of testing. These authors also showed that NMDA receptors in DG granule cells are important for context-specific modulation of the firing rate in CA3. Surprisingly, mice lacking the α5-subunit of the GABA_A_ receptor in DG granule cells also show a rapid pattern separation deficit, in that the discrimination ratio is different starting on the second day of testing^21^. Thus, two mouse models modulating the activity of DG granule cells, one likely inhibiting them (by ablating an excitatory receptor) and one potentially activating them (by attenuating tonic inhibition) disrupt rapid pattern separation. These conditional knockout experiments thus suggest that both activation and inactivation of DG granule cells may disrupt rapid pattern separation, suggesting that the normal activation level of DG granule cells may be optimal for this task. Ablating young adult-born DG granule cells has also been reported to reduce rapid pattern separation capability^23^. Enhanced rapid pattern separation was reported in triple transgenic mice in which the output of old DG granule cells was inhibited^23^. Interestingly, these mice also exhibited deficits in rapid pattern completion^23^ which we have also observed in the CalEx mice. One interpretation is that inhibition of astrocytes would indirectly inhibit old DG granule cells, which would be consistent with the lower average intensity of DG neuronal calcium activity we observed (Fig. 10).

From our current data alone, the precise mechanisms by which this inhibition of DG neurons occurs are unclear. However, a recent report examining the role of GABA transporter 3 (GAT-3) in DG astrocytes has provided some insight into a potential mechanism. When CalEx was expressed in DG astrocytes, a GABA-induced increase of mEPSC frequency in DG granule cells is no longer observed^38^. Together with other findings, the study concluded that GABA released by interneurons is taken up via GAT-3 by astrocytes, which induces influx of calcium into the astrocytes. Such calcium influx then results in a release of glutamate from the astrocytes, which acts on presynaptic GluN2B receptors on neuronal projections of the perforant pathway, resulting in enhanced glutamatergic synaptic transmission onto DG granule cells. Thus, disruption of astrocytic calcium influx by expression of CalEx would be expected to effectively inhibit DG granule cells. This is consistent with our observation of a lower average neuronal calcium activity in the DG neurons of CalEx mice. Taken together, our findings and the other study establish a plausible mechanism by which CalEx expression in DG astrocytes results in the decreased activity of DG neurons that we observed. The other study also reported that mice expressing CalEx in the DG showed reduced contextual fear conditioning^38^. Although our contextual pattern separation experiment uses different fear conditioning training protocols compared to standalone fear conditioning protocols, during the training period we did observe some evidence that may indicate that CalEx mice have a marginal deficit in contextual fear conditioning (Fig. S5). It is notable that apart from rapid pattern separation, enhanced delayed or learned pattern separation has been described in global GABA_A_ receptor α5-subunit knockout mice. Global GABA_A_ receptor α5-subunit knockout mice and CA1 GABA_A_ receptor α5-subunit knockout mice both have similarly reduced discrimination ratio compared to α5-floxed controls on days 1 and 2, but the global α5 knockout mice discriminated the two contexts better than both α5-floxed control mice and CA1 α5 knockout mice on days 9 and 12^36^. In our current study we observed that while inactivation of astrocytes via CalEx expression not only improved rapid pattern separation but also learned pattern separation, activation of the G_q_ GPCR hM3Dq only improved learned pattern separation, but not rapid pattern separation. These results indicate that astrocytes differentially regulate these two forms of pattern separation.

Since GPCR DREADDs were developed earlier than CalEx, so far, more studies have utilized these tools and shown their effects. Studies of astrocytes in other brain regions like CA1 ^10,14,15^ based on activation of GPCR DREADDs have linked astrocytic function to the neuronal function of the respective local brain region. Indeed, we observed an increased contextual pattern separation learning rate in G_q_ DREADD mice and a disruption of different types of fear extinction (within-session and between-session fear extinction) in a pathway-specific manner. Although both G_i_ and G_q_ GPCRs in astrocytes have been shown to trigger calcium-dependent release of gliotransmitters like glutamate and D-serine^10,39,40^, previous studies in other brain regions have also shown that activation of the G_i_ and G_q_ GPCR DREADDs in astrocytes may have different behavioral outcomes^10,16^, which is consistent with our observations. Since G_i_ and G_q_ DREADD activation used in these studies are predicted to induce downstream calcium signaling^6^, it was expected that while the observations from the G_q_ DREADD and the G_i_ DREADD activations might differ, at least one of the GPCR DREADDs would have an effect on behavioral phenotypes opposite to that of CalEx. However, in G_q_ DREADD mice or G_i_ DREADD mice we did not observe behavioral changes opposite to those seen in CalEx mice.

This indicates that the astrocytes’ integration into the neuronal networks of the DG is highly complex and that our observations in CalEx mice are likely an integrated response from multiple affected (G_q_-dependent and G_i_-dependent) pathways that each individually may enhance or inhibit neuronal function. Additionally, although the G_i_ DREADD is expected to induce downstream calcium-dependent signaling, the actual observed effects on calcium signaling depend on the brain region ^14,41^. Thus for the DG it is still unclear whether chronic G_i_ DREADD activation actually increases astrocytic calcium dynamics. In general, it is important to view expression of CalEx, chronic G_i_ DREADD activation and chronic G_q_ DREADD activation as three distinct methods of manipulation that may affect overlapping pathways and functions. Within-session fear extinction has previously been found to be facilitated by CB1 receptor signaling^31^ which is needed for spike timing-dependent long-term depression (t-LTD) at granule cell synapses with inputs from the perforant pathway^42^. This process requires glutamate released by astrocytes^42^ which can be triggered by astrocytic calcium^43^. However, astrocytic CB1 receptors in other brain regions have also been shown to regulate long-term potentiation (LTP) by releasing ATP and D-serine^44,45^. Thus, our chronic G_q_ DREADD activation may have disrupted both t-LTD and LTP which are facilitated by the CB1 receptor pathway. NMDA receptors have been shown to facilitate between-session fear extinction^46^. As mentioned previously, G_i_ GPCR pathways are involved in astrocytic release of glutamate^39,40^ which potentially indicates that chronic G_i_ DREADD activation disrupted normal NMDAR function in DG.

As outlined above, we hypothesize that astrocytes in the DG promote the activity of old granule cells, and our observations made in CalEx mice support this hypothesis. However, also as mentioned above, likely more than one pathway or mechanism is involved in the astrocytic regulation of calcium activity in DG neurons. Indeed, the observation of an increased calcium event rate in CalEx mice may indicate that there is a mechanism at work which is weakly excitatory for neurons. Previous studies in the striatum have found that inhibition of calcium signaling with CalEx increases expression and protein level of GABA transporter 3 (GAT-3), reducing tonic GABA currents and causing repetitive behavior which can be rescued by blocking GAT-3^47^. Moreover, spike timing-dependent long-term depression at entorhinal cortex to granule cell synapses is dependent on astrocytic calcium^42^, and might thus be disrupted by expression of CalEx. Both increased GAT-3 expression and disruption of t-LTD are expected to be overall excitatory for DG neurons. In addition to disruption of the previously mentioned calcium-dependent release of glutamate and D-serine, it is thus conceivable that CalEx might elicit both excitatory and inhibitory effects on the activity of the dentate gyrus neurons, the integration of which may underlie its complex effects on the calcium signal and behavior we have observed.

Our study has limitations that need to be taken into account when interpreting our results. Our fear extinction experiments have been conducted after a contextual pattern separation paradigm with a high number of foot shocks before starting the extinction training. It will be interesting to see whether the same types of observations will be made using a more standard fear conditioning and fear extinction protocol. While we were doing calcium recordings in the home cage, the mice showed no overt signs of abnormal behaviors or seizures. Later on, some of the mice developed seizures and showed degraded nesting behavior (Fig. S6) which has been reported by other studies investigating mouse models with seizures^48,49^. The GCaMP6f concentration used in our study has been shown in a previous study of the effects of GCaMP on the dentate gyrus to be insufficient to cause seizures^50^. However, it is conceivable that the combination of hippocampal damage induced by the procedure of lens implantation and GCaMP expression may have contributed to the development of seizures in control mice. It is notable that in CalEx mice more cases of seizures were observed than control mice (Fig. 2), especially among the mice in which we were able to record calcium signals which is an indication that the GRIN lens was placed close to the DG. While our sample size is limited, a preliminary statistical test shows that among the animals for which calcium signals were recorded, CalEx mice do show a higher chance of developing seizures (Chi-square test, p=0.0209). Thus, it could be argued that expression of CalEx in DG astrocytes may increase the likelihood of developing seizures for mice with preexisting surgical lesions and expressing GCaMP6f. It is noteworthy that we did not observe seizures in any other CalEx mice that had no lens implantations.

Overall, our findings further confirm the significant role of astrocytes in neural circuit function in the dentate gyrus and help to fill the knowledge gap in understanding the behavioral significance of DG astrocytes. Future studies could explore whether a change in the activity of DG astrocytes also has effects on neuronal circuits outside of the DG. A deeper understanding of how astrocytes contribute to the integration of interconnected neural circuits will enable the development of novel therapeutic strategies targeting astrocytes.

## Acknowledgements

This work was funded by the University of Illinois Urbana-Champaign. The authors declare no competing interests.

