## Supplemental Figures S1-S6 for "Astrocytes in the Dentate Gyrus Shift the Equilibrium from Pattern Separation to Pattern Completion and Inhibit Fear Extinction"

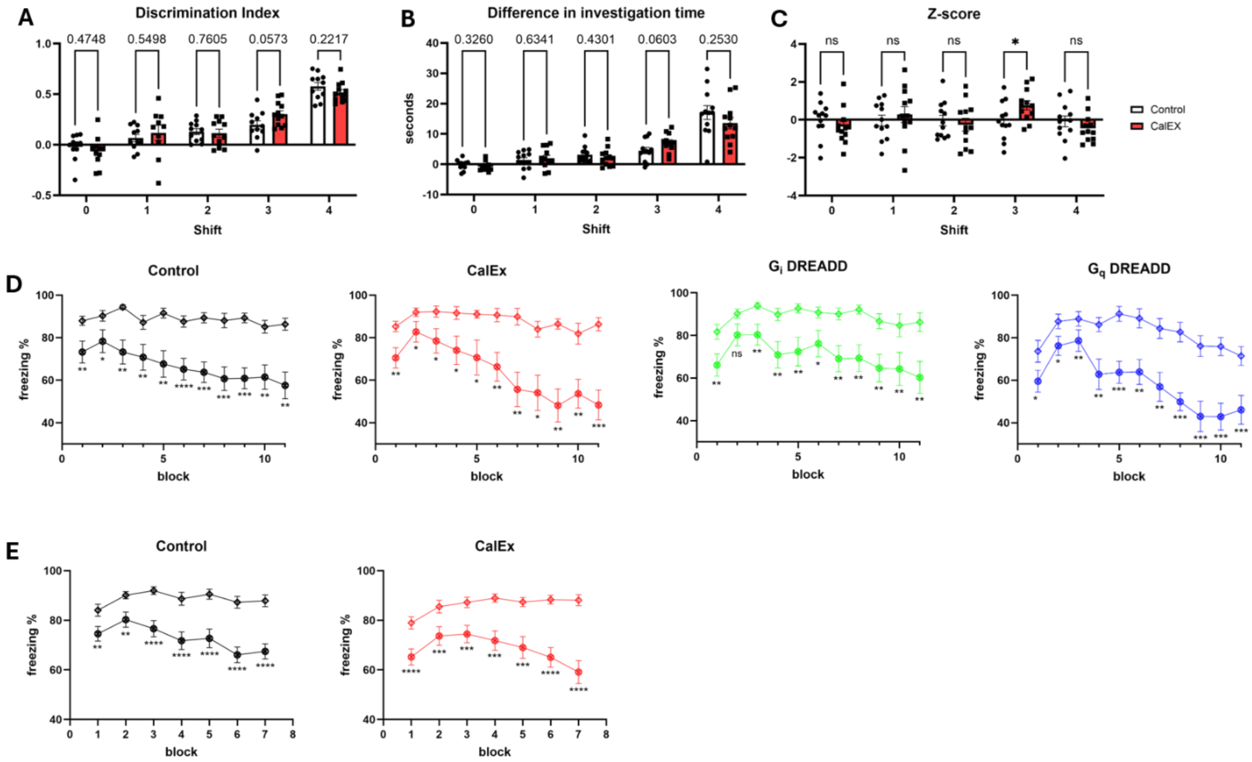

**Fig. S1 Analysis of object pattern separation and contextual pattern separation.** CalEx mice showed trend of improved **(A)** discrimination index and **(B)** difference in investigation time at shift distance 3 in object pattern separation experiment. **(C)** z-score analysis of the parameter in (**A**-**B**) showing CalEx mice have improved performance at shift distance 3. **(D)** Freezing percentage in each of the two contexts from contextual pattern separation protocol for animals in batch 4. **(E)** Freezing percentage in each of the two contexts from contextual pattern separation protocol for batch 3 and batch 4 combined. Two-way ANOVA repeated measures was used for all panels, n=22, n=11 for male and female sub-population, *p < 0.05, **p < 0.01. Data presented as mean ± SEM.

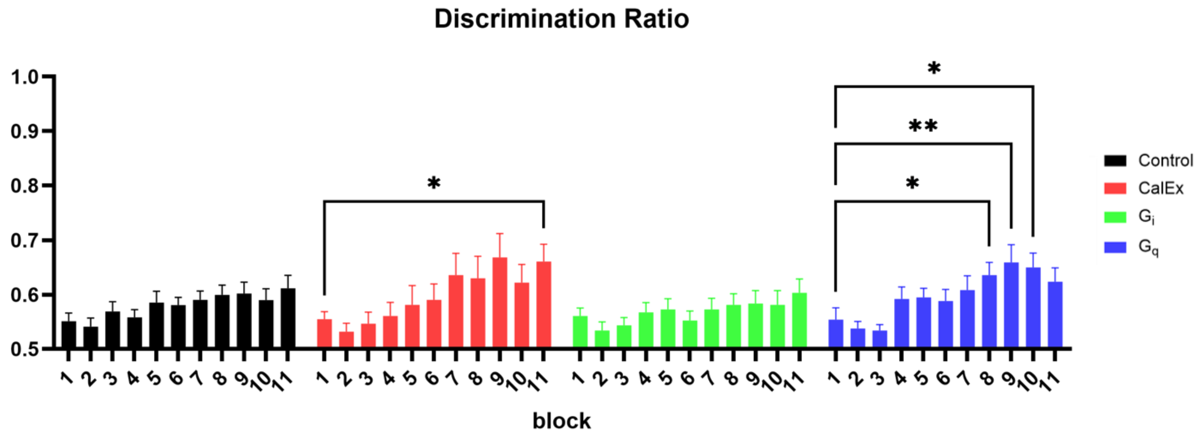


**Fig. S2 CalEx and $G_q$DREADD potentially have accelerated learning rate in contextual pattern separation.** CalEx mice and $G_q$ DREADD mice show significant increase in discrimination ratio at later blocks compared to blocks 1. Data from this panel originates from behavioral batch 4. Two-way ANOVA repeated measures, n=10, *$p < 0.05$, **$p < 0.01$. Data presented as mean ± SEM.

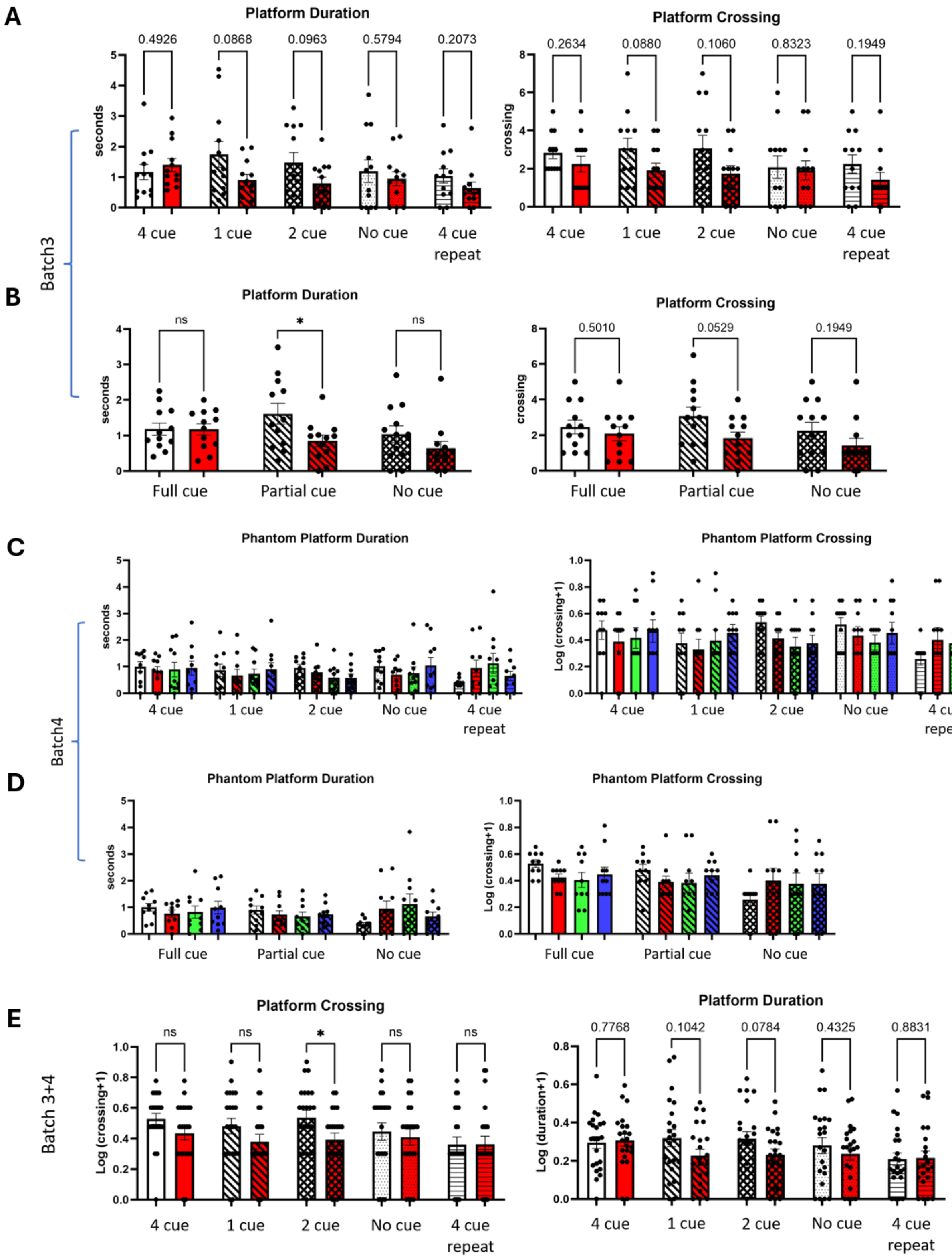


**Fig. S3 Analysis of pattern completion experiment separated by batches.** Phantom platform duration and crossing count analysis for batch 3 analyzed by individual cue conditions **(A)** and by combined cue conditions **(B)**. (**C**-**D**) Similar analysis for batch 4. (**E**) Phantom platform duration and crossing count analysis for batch 3 and batch 4 combined analyzed by individual cue condition. Two-way ANOVA repeated measures, n=11-12 for (**A**, **B)**, n=10 for (**C**, **D**), n=23-24 for (**E**), $*p < 0.05$. Data presented as mean ± SEM.

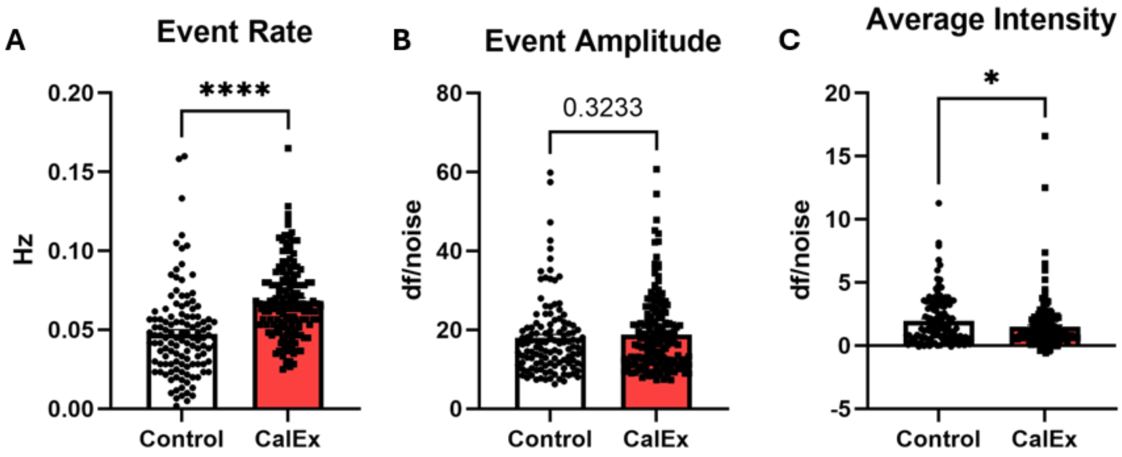


**Fig. S4 Analysis of in-vivo calcium imaging including the mice with lens placement marked as "likely".** Comparison of neuronal calcium event rate **(A)**, event amplitude **(B)** and average signal intensity **(C)** between control and CalEx mice. Mann-Whitney test, n=121-167. *$p < 0.05$, ****$p < 0.0001$. Data presented as mean ± SEM.

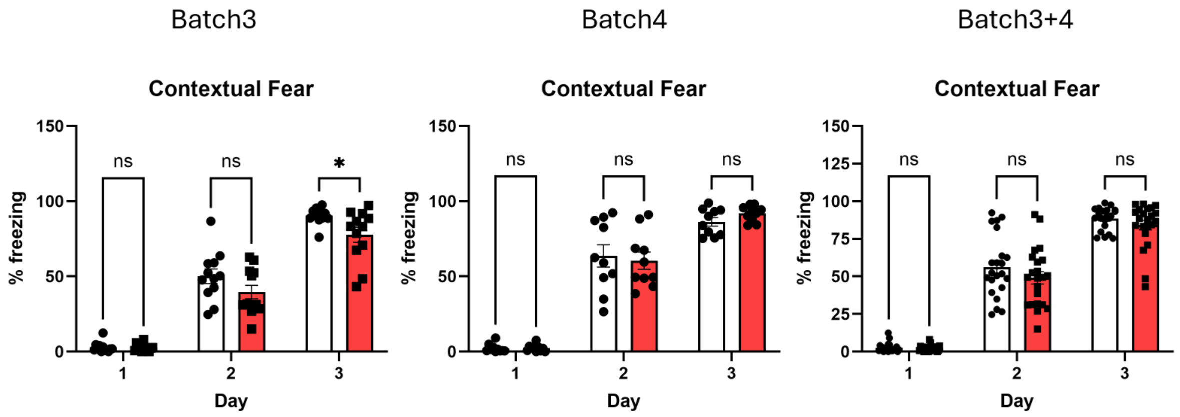


**Fig. S5 Analysis of freezing percentage during the first 3 days of contextual fear conditioning of contextual pattern separation experiment.** CalEx mice in batch 3 showed reduced freezing percentage on day 3 of contextual fear conditioning compared to control mice. No difference was observed for batch 4 and batch 3 and batch 4 combined. Two-way ANOVA repeated measures, n=11-12 for batch 3, n=10 for batch 4, n=23-24 for batch 3 and batch 4 combined, $*p < 0.05$. Data presented as mean ± SEM.

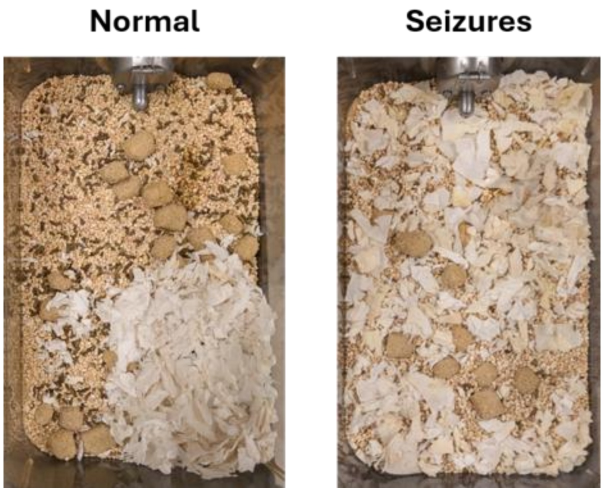


**Fig. S6 Example of degraded nesting behavior for mice showing seizures.** Representative example of nesting in home cage for normal mice and mice that developed seizures.
